# Generating synthetic aging trajectories with a weighted network model using cross-sectional data

**DOI:** 10.1101/2020.02.14.949560

**Authors:** Spencer Farrell, Arnold Mitnitski, Kenneth Rockwood, Andrew Rutenberg

## Abstract

We develop a computational model of human aging that generates individual health trajectories with a set of observed health attributes. Our model consists of a network of interacting health attributes that stochastically damage with age to form health deficits, leading to eventual mortality. We train and test the model for two different cross-sectional observational aging studies that include simple binarized clinical indicators of health. In both studies, we find that cohorts of simulated individuals generated from the model resemble the observed cross-sectional data in both health characteristics and mortality. We can generate large numbers of synthetic individual aging trajectories with our weighted network model. Predicted average health trajectories and survival probabilities agree well with the observed data.

## Introduction

Human aging is a complex process of stochastic accumulation of damage^1^ that occurs at many organismal scales ranging from the cellular^2^ to the functional. Individual health trajectories are heterogeneous, but typically worsen with age as damage accumulates. Heterogeneity of aging trajectories arises even in studies of clonal organisms in controlled laboratory conditions,^3, 4^ and is an intrinsic part of aging. Heterogeneity in health as individuals age has been measured with a variety of methods, although here we focus on binary “health deficits” determined from routine clinical assessment and self-reported surveys.^5–9^ Health deficits are indicators of an aging phenotype, indicating disease, laboratory abnormalities, cognitive impairment, disability, or difficulty performing everyday tasks.

While any single deficit may not be a good measure of overall health, or a very informative predictor of mortality, averaging many binary deficits to evaluate overall health provides measures that are strongly associated with both adverse health outcomes and mortality.^5–10^ Furthermore, such frailty indices (FIs) are robust to missing or heterogeneous data.^8^ Using “high-level” health-deficits provides a measure of health that is both conveniently assessed and reflecting the functional aspects of healthy living that are important to the individual.^11^ Such an FI also contains information about health that is not found in recent epigenetic measures based on DNA-methylation.^12^

While the FI has been shown to be broadly predictive of both mortality^13^ and of the accumulation of individual deficits,^9, 14^ it does not distinguish the health trajectories of two individuals with the same FI even if they have distinct sets of accumulated deficits. Capturing the heterogeneity in health trajectories requires modelling the full high-dimensional set of health variables. We develop a model to generate populations of synthetic health trajectories, which capture this heterogeneity.

However, the development of models of aging is complicated by the data currently available. Large observational studies (10^4^+ individuals) with linked mortality are often cross-sectional (measuring most variables only when individuals enter a study), have short censored survival outcomes, and have a lot of missing data. This has made developing realistic models of human aging difficult. Nevertheless, such models would be useful to generate model individual health trajectories during aging from birth, or from baseline data of actual individuals. Encouragingly, a model capable of generating general health trajectories during aging using cross-sectional data has recently been developed^15^ – though it did not consider individual survival.

Here, we develop an intuitive model that can be fit with cross-sectional data with censored survival information to generate individual aging trajectories that include both health *and* survival. Our model is adapted from previous work modeling human aging with stochastic dynamics on a complex network,^16–18^ which was shown to capture population level aging phenomena, such as Gompertz’ law of mortality.^19^ This model was based on the intuitive assumption that having one health deficit increases the risk of acquiring another one, and so deficits can be thought of as interacting in a network, in which connections establish pairwise associations.^20^ Nodes in this older model represented generic/abstract deficits and corresponded to no specific physiological systems in particular. Nevertheless, their collective behaviour captured key aspects of aging. This “generic” network model (GNM) used many nodes (*N* = 10^4^) to abstractly represent the many interacting physiological systems in the human body, had simple interactions between nodes, and needed no age-dependent programming of damage.

Our new “weighted” network model (WNM) is parameterized so that each node represents an actual health attribute (potential health deficit) corresponding to observational aging data. We recognize that we will never be able to incorporate all possible health attributes as nodes in our network or to describe exact biological mechanisms. For this reason, we use more complex weighted interactions between observed nodes that can capture the effective behaviour of underlying and/or unobserved biological mechanisms. This new WNM can be considered as a coarse-grained adaptation of our previous GNM, with far fewer nodes.

We separately fit our WNM with cross-sectional observational data from the Canadian Study of Health and Aging (CSHA)^21^ and the National Health and Nutrition Examination Survey (NHANES).^22^ These human aging studies consist of 8547 and 9504 individuals, age ranges of 65 − 99 and 20 − 85 years, in which mortality data are available for at most 6 or 10 years past study entry, respectively. Deficits in these datasets are binary indicators of health issues and integrate information across physiological systems, such as difficulty performing activities of daily living (ADLs) or more complex instrumental activities of daily living (IADLs). We estimate parameters for each study by maximizing the log-likelihood for our model to recover the observations, where the likelihood is estimated from simulations of the stochastic model. By validating our model on a separate test set, we demonstrate that our model represents real aspects of aging, and is not overfitting to the training data.

We find that our synthetic individuals generated with our model capture the health outcomes and survival of observed data with a number of different measures. Indeed, rather than focusing on achieving optimal predictive performance on any one particular task, our goal was to obtain a robust model that can generate realistic trajectories for multiple health attributes at once for many individuals from either actual or synthetic baseline health status. Nevertheless, our model cannot overcome the intrinsic limitations of cross-sectional data –– for example, the accuracy of health trajectories will only be assessed by comparing simulated cohorts of individuals to longitudinal data of the observed cohort.

## Results

### Health trajectories

Starting from an individual with a set of *N* potential binary deficits at a baseline age *t*_0_, 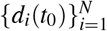, our model (see Model section below, trained on the CSHA dataset) generates deficit trajectories {*d*_*i*_(*t*)} describing health for synthetic individuals for each age *t* > *t*_0_ until mortality. (Here *d*_*i*_ = 1 indicates a deficit for the *i*th aspect of individual health, while *d*_*i*_ = 0 indicates no deficit. We generally refer to a set of potential health deficits {*d*_*i*_} as health attributes). We want to test whether these synthetic individuals age with the same properties as do real individuals in the observed data. Without longitudinal data, we cannot test these individual trajectories directly. However, we can use the population average of the observed cross-sectional data and compare with the average population trajectory predicted from our model. If the study population was randomly sampled with no biases, we expect these average trajectories to agree.

Given baseline age and *N* = 10 selected deficits for individuals from the test data aged 65 - 70 that survive past age 70, and individuals aged 75 - 80 that survive past age 80, we use our model to forecast their health trajectories. These simulated trajectories allow us to the compute the full joint distribution of health states vs age for these simulated individuals, 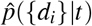. Since this distribution is 10-dimensional, with 2^10^ = 1024 distinct age-dependent probabilities, we plot only the prevalences of each deficit (by marginalizing the distribution). We compare the average of these individual trajectories until death to the deficit prevalence from the observed cross-sectional CSHA data for ages 70+ and 80+. In Figure 1 we show health trajectories using prevalence at age *t* of the *i*th deficit, 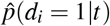, for the model (blue solid lines for 65 - 70 and green dashed lines for 75 - 80) together with the observed CSHA prevalence (red squares). We see excellent agreement for nearly 30 years for most deficits. This shows that the model is able to project a population forward in time while correctly identifying changes in deficit prevalence. We also show the average predicted trajectories with an alternative set of 10 deficits in Supplemental Figure S4 and using the NHANES data in Figure S5.

**Figure 1.**
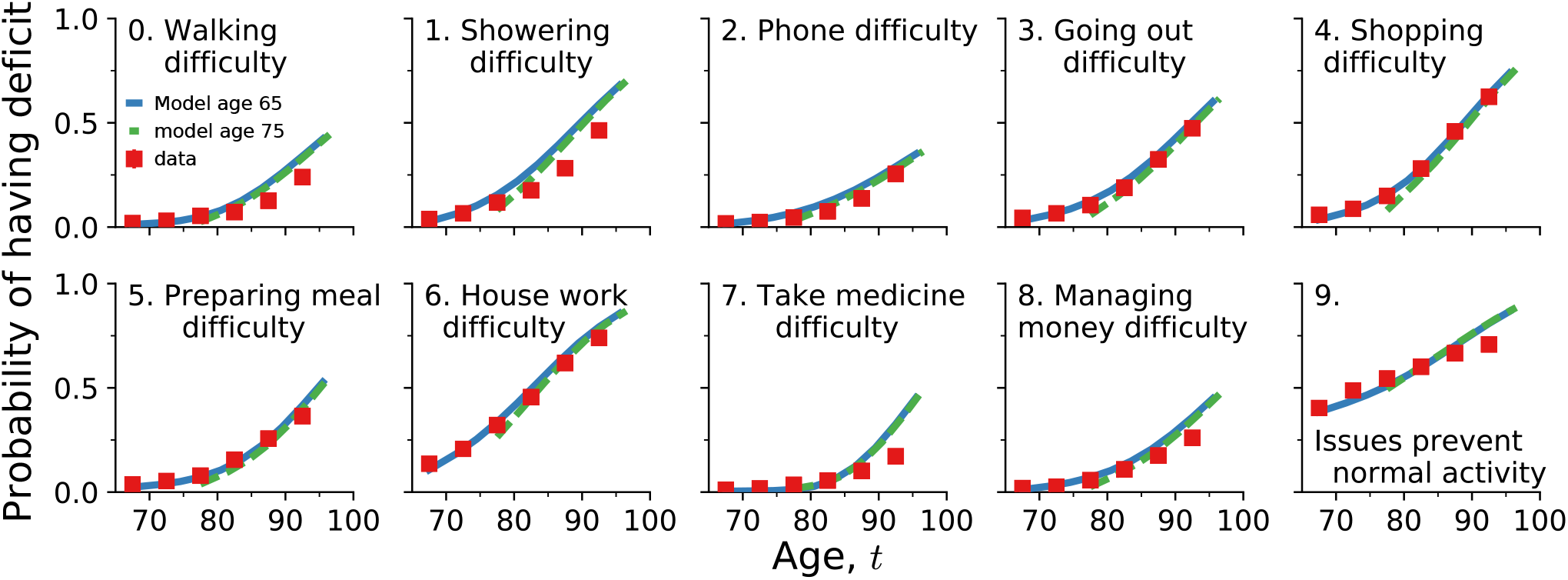
Average predicted trajectories of ten deficit prevalences (as indicated by the subplot titles, numbered 0 to 9) vs. age for individuals from the test data aged 65 – 70 surviving past 70 (solid blue lines) and aged 75 – 80 surviving past 80 (dashed green lines). Observed CSHA prevalence is shown in red squares; standard errors are smaller than the point size.

A similar prediction is done for the prevalence of pair combinations of deficits, i.e. comorbidities. We predict the probability of having two specific deficits 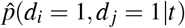 vs age in our model, and compare with the observed data. This is shown in Figure 2 and in Supplemental Figure S6, and also shows excellent agreement between the model and the CSHA data for over 30 years. This indicates that our model is accurately capturing the association between pairs of deficits through network interactions. Notably, pairwise combinations in the model often perform better than corresponding prevalences – see for example (1, 7) in Figure 2 – confirming that pairwise combinations (together with higher order combinations, not shown here) are non-trivial predictions of the model.

**Figure 2.**
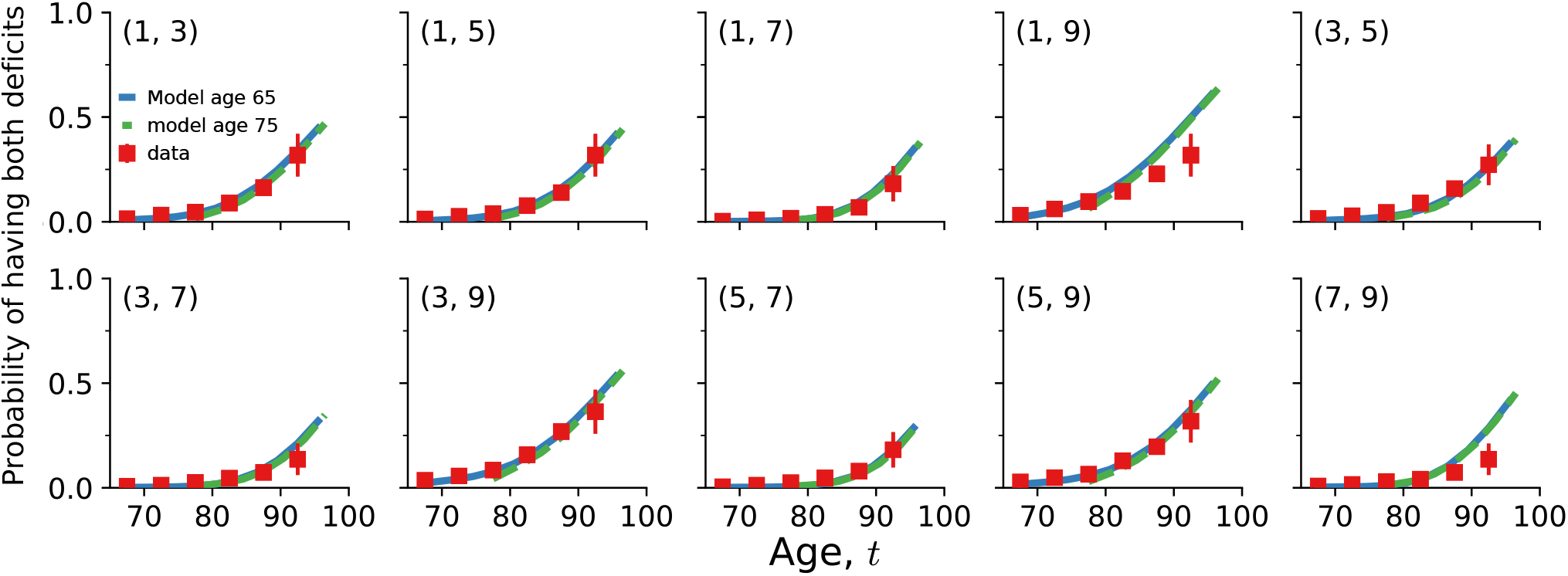
Average predicted trajectories of pairwise deficit prevalence vs. age for individuals from the test data aged 65 – 70 surviving past 70 (blue solid lines) and aged 75 – 80 surviving past 80 (dashed green lines). Subplot titles indicate the two deficits included in the pairwise prevalence, with numbers corresponding to titles in Figure 1. Only odd numbered pairs are shown here; other pairs are shown in Supplemental Figure S6. CSHA data is shown by red squares; errorbars represent standard errors of the pairwise prevalence.

We can represent overall individual health with the well-established “Frailty Index” (FI), an index that uses the proportion of deficits, 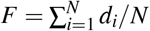, as a predictor of health and mortality.^5, 8^ In Supplemental Figure S7 we show that the heterogeneity in health as individuals age, as characterized by distributions of FI at different ages, is similar to the observed CSHA data.

We are not limited to modelling known individuals. In Supplemental Figure S8 we show that the population prevalences of synthetic individuals starting from birth with zero damage also agree with the observed CSHA prevalences. Indeed, we can generate trajectories and survival curves for any baseline age and individual set of deficits. Using synthetic populations, we can also generate trajectories starting from any age with partially observed sets of deficits with missing values.

Figure 3 shows FI trajectories starting from the known baseline data (red circle) for 6 synthetic individuals with specific deficits. Horizontally, we vary baseline age with 65, 75, and 85 along the columns. Vertically, we vary baseline deficits, with bottom individuals having a higher initial FI by having two additional deficits. Individual trajectories are conditioned on dying at their median survival probability (dashed black lines), seen from the individual black survival curves. Shaded regions show a distribution of FI trajectories. The trajectories behave reasonably. Individuals with more baseline deficits accumulate additional deficits faster and die sooner. Individuals starting at older ages also have a more rapid increase in number of deficits and have a shorter time to death. Note that these trajectories exhibit FIs larger than the typically observed maximum of 0.7,^9, 23, 24^ which is due to the small number of potential deficits used here (only 10) compared to typical studies (with 30 - 40 potential deficits).

**Figure 3.**
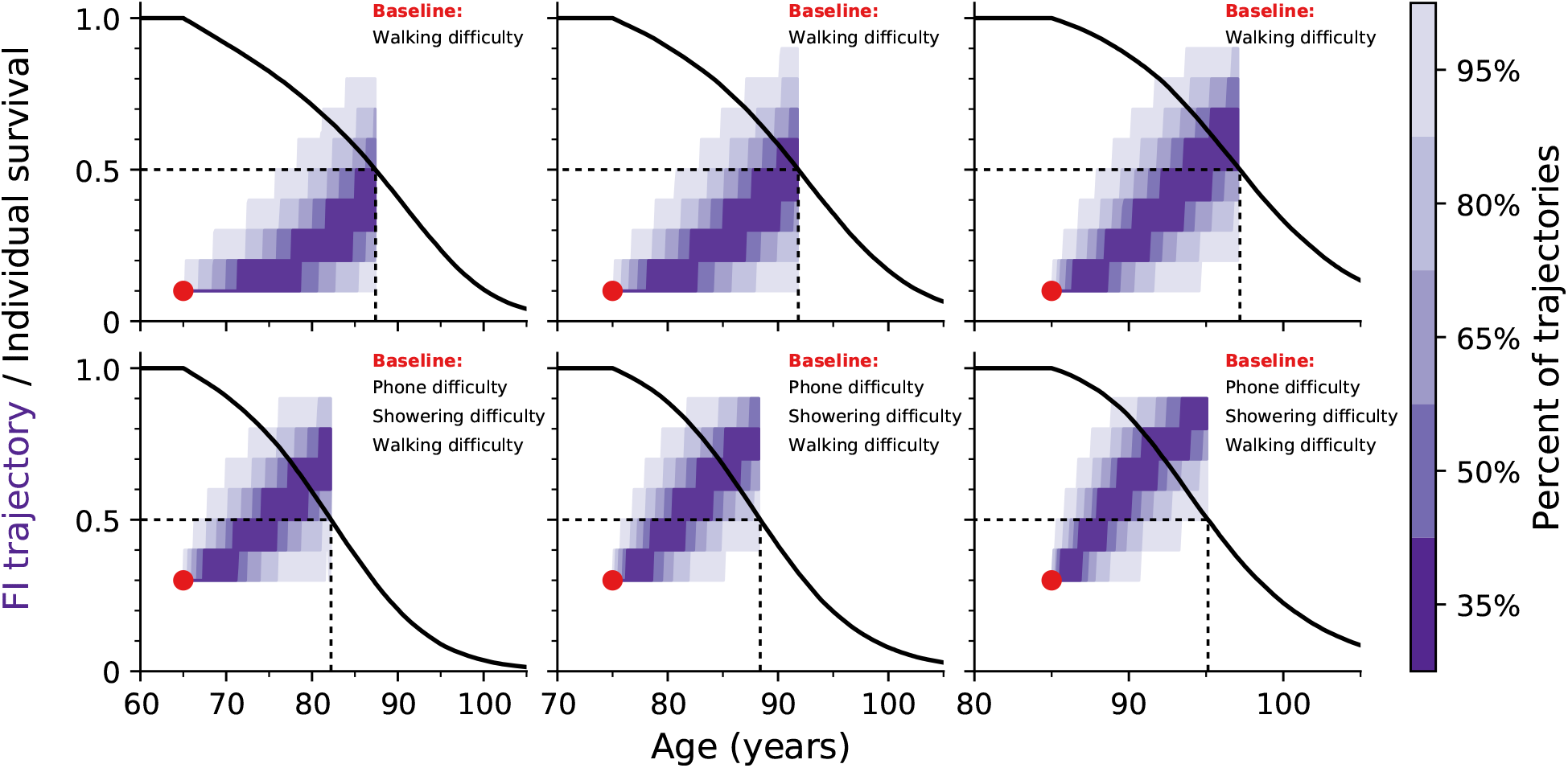
The purple shading indicates the distribution of Frailty Index trajectories vs age for simulated individuals starting from the red circle at a specific age, with the indicated starting set of deficits. The scale-bar at right indicates the probability that a simulated trajectory falls within a particular colour. In the top row individuals only start with one deficit. In the bottom row individuals start with three deficits. In the three columns, individuals start at age 65, 75, or 85. Individual survival curves are shown as black lines, and trajectories are conditioned on dying at the median death age (indicated with dashed lines).

### Individual deficit predictions

Since our training and test data sets have similar distribution of health states (e.g. deficit prevalence), the good test performances in Figure 1 and Figure 2 do not rule out overfitting to the training set, because the model was only assessed against the distribution of health states for the training and test data. To assess overfitting, we need to consider predictions of individual health states – i.e. observed deficits for specific individuals. We first verify that there is only a ~ 20% overlap between the observed health states of individuals in the training and test data sets, see Supplemental Figure S9. (We also confirm that using only one or two attributes leads to a ≳ 90% overlap. Note that this overlap is not due to the same individuals being present in the training and test sets, but due to the discrete nature of the binary deficits.)

We test the model’s ability to capture the age-dependent joint distribution of deficits *p*({*d*_*i*_}|*t*) by evaluating its performance in predicting “left out” deficits from individuals in the test set at the same age, i.e. performing missing data imputation by estimating *p*({*d*_*j*_}_missing_|{*d*_*i*_}_observed_, *t*). Given a known age *t*^(*m*)^ and known health attributes {*d*_*i*_}^(*m*)^ for an individual *m* from the test set, we simulate from zero damage at birth and sample from the simulated individuals at age *t*^(*m*)^ that have the health attributes 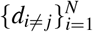 to estimate the probability of having the left out deficit, 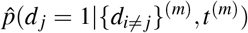. We compare this probability with the actual value of the left out health attributes 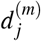 for this individual at the same age. This is a binary classification, and can be quantified by the area under a ROC curve – the AUC.

Figure 4 shows the AUC for each left-out deficit. This is equal to the probability that given two individuals with and without the deficit, the model correctly predicts a higher probability of having the deficit for the individual with the deficit than without. The full test set is shown as a solid blue line, while blue circles show the AUC stratified by age. For all deficits, we see AUC values well above 0.5 for the test data, meaning that our model is making informative predictions and not just overfitting the training data. Orange squares and lines show similar values for the AUC of the training data, which is further confirmation that any overfitting is minimal.

**Figure 4.**
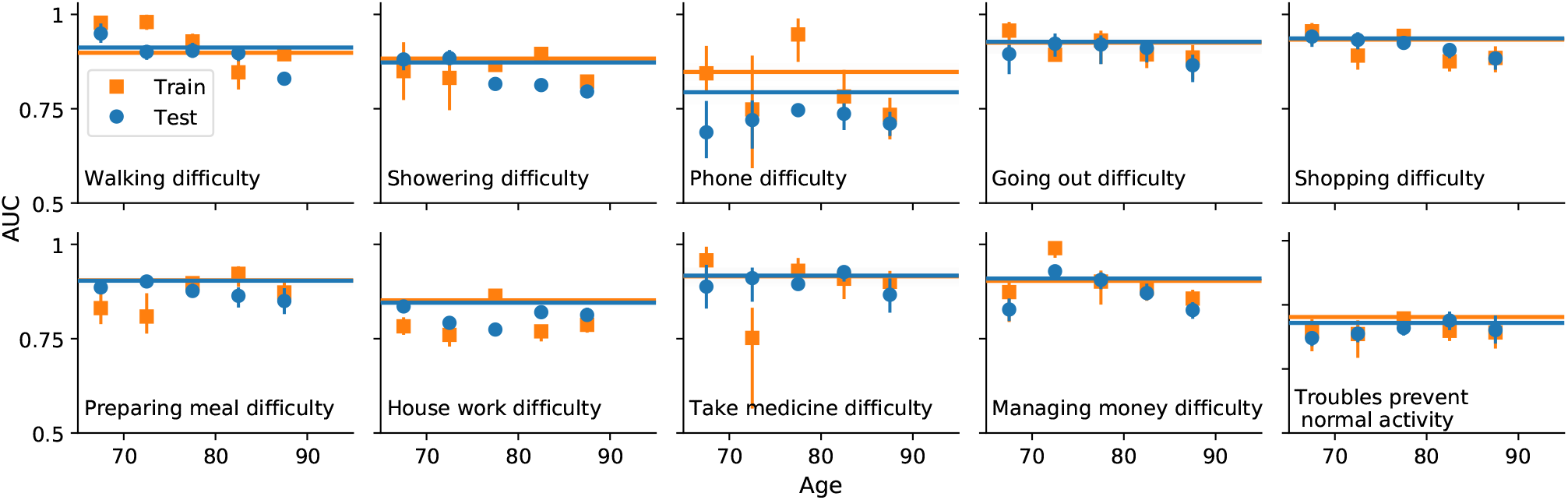
AUC for predicting the value of individual left-out deficits for test (blue circles and lines) and training (orange squares and lines) data. The points show the AUC when stratifying by age. Errorbars indicate 95% confidence intervals. The solid line shows the result for all ages. The baseline AUC is 0.5 for random (uninformative) predictions.

Obtaining essentially the same AUC when stratifying by age in Figure 4 demonstrates that the model is predicting as well even when we eliminate age as a factor. This indicates that the model is utilizing the network interactions between observed deficits to make non-trivial individual predictions, and not just using e.g. increasing prevalence with age.

### Survival

We can also model individual mortality. We take baseline data from the test set for an individual *m* at age *t*^(*m*)^ and health attributes {*d*_*i*_}^(*m*)^, and simulate from this age until mortality. This allows us to estimate their survival function 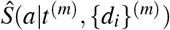, i.e. the individual’s probability of surviving to an age *a* > *t*^(*m*)^. We average these to get a population average survival function over *M* individuals, 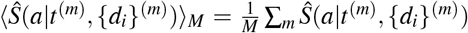. We show the comparison of this to a Kaplan-Meier estimate of the population survival function from the observed CSHA test data in Figure 5A, with our model shown in blue and the observational test data in red. We observe good agreement, and also find that model predictions correctly drop to zero survival by age 120.

**Figure 5.**
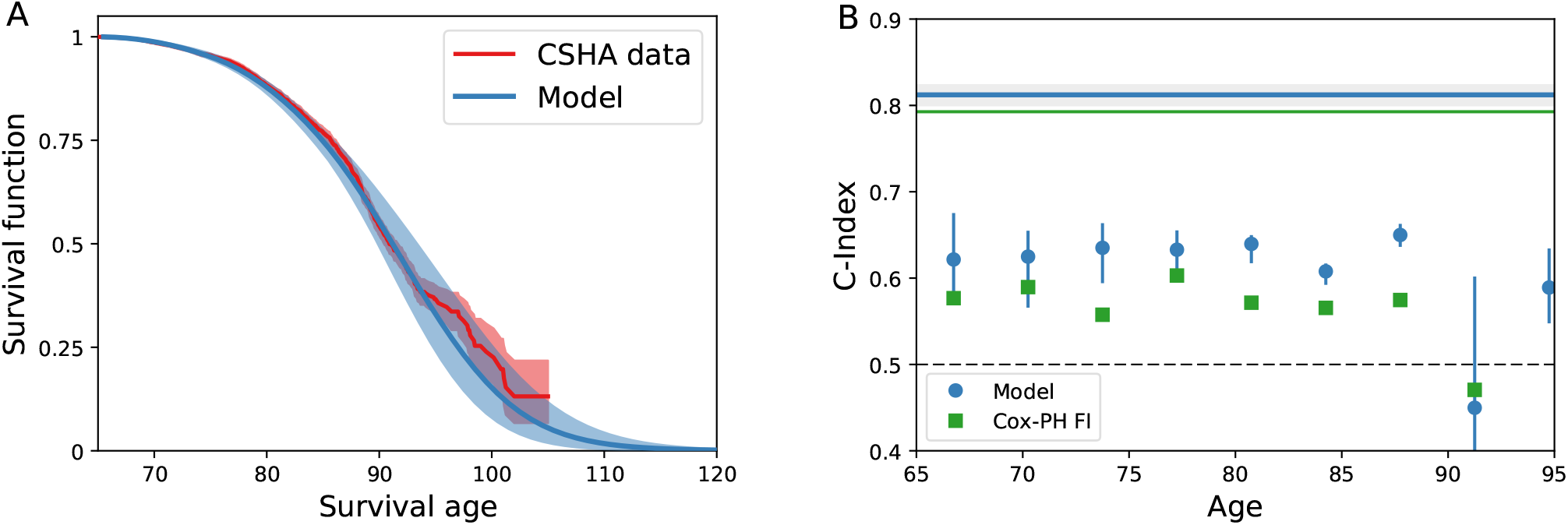
A) Model population average survival function (blue) and CSHA Kaplan-Meier population survival function (red). Error in the model and data are shown as shaded regions for a 95% confidence interval. B) Survival C-Index stratified by age (blue circles) and unstratified (solid blue line). Green squares and lines show the age-dependent *C*^td^ for a Cox-proportional hazards model with age and FI. Errorbars on points and grey region around the unstratified line show 95% confidence intervals.

Since the training and test distributions are similar, a population measure of mortality does not tell us whether the model is overfitting – only whether the model is able to capture the population trends in mortality. Accordingly, we validate individual survival on the test set with a C-index^26^ to measure how well the model discriminates individuals in terms of risk of mortality. The C-index is the probability that the model correctly predicts which of a pair of individuals lives longer, so a C-index above is making predictions better than random.

Since our model includes potentially complex time-dependent effects, where survival curves can potentially cross, we use a more general age-dependent C-index.^27^ We obtain this by comparing the rank ordering between survival probability and known survival age while including censoring, so 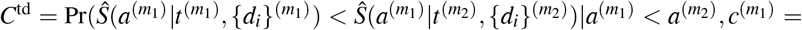 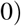.^27^

Figure 5B shows this age-dependent C-index for both on the full test set (solid blue line) and stratified by age (blue circles). The C-index shows that the model discriminates well on the full test when the difference in ages between individuals can be used in the discrimination. When we stratify by age to eliminate this effect, we nevertheless see that the model still discriminates well based on just these 10 deficits alone, indicating that the model captures an increased risk of mortality from specific deficits. In particular, our model performs better than a standard Cox-proportional hazards^28^ model using the Frailty Index and age (green squares and line). We note that stratified values are noisy due to the small number of individuals per age bin, especially at higher ages.

In Supplemental Figure S10A we show similar results for the C-Index for both training and test sets, which also indicates a lack of overfitting. Similarly, Figure S10B shows an *R*^2^ measure constructed from Brier Scores,^29^ a measure of how well predicted and observed survival curves match, that behaves similarly for training and test data. Furthermore, Figure S10C shows the ROC AUC for predicting binary dead/alive on the train/test sets within a specific window of time, finding a similar AUC of approximately for 1-5 year mortality windows. For all of these, we find similar behavior between the training and test sets, indicating a lack of overfitting. This is also seen for survival predictions for the alternative set of deficits used in Figure S4, as shown in Figure S11 and for survival predictions for the NHANES dataset, as shown in Figure S12.

## Discussion

Our weighted network model (WNM) is trained with cross-sectional data, generates cohorts of synthetic individuals that resemble the observational data, and can forecast the future survival of real individuals from their baseline health and age. We have validated the WNM model through a variety of measures. Synthetic individuals age with trajectories that have approximately the same prevalence of deficits and comorbidities as in the observed data. The average trajectories predicted by the model agree very well for nearly 30 years. Given a set of known deficits, the model can predict the probability of having a missing or unknown deficit at the same age, demonstrating the models ability to capture the age-dependent joint distribution of the deficits. Estimated survival curves also agree with observed population survival, are predictive of mortality, and discriminate between individuals.

Our model has a large number (188) parameters while our health data has only *N* = 10 binary attributes. One concern with having at most 2^*N*^ = 1024 discrete health states is that there could be significant overlap between the test and training sets. Nevertheless, we showed in Supplemental Figure S9, that a significant fraction of the health states from the test set are not in the training set. Using this (non-overlapping) test set we have shown that our model does not substantially overfit and can make predictions on unseen test individuals.

We emphasize that our model is not just fitting the prevalence of the *N* = 10 deficits, but is fit to the full 10-dimensional age-dependent joint distribution *p*({*d*_*i*_}|*t*) which has 1024 distinct states. A full model of this distribution would require many more than 1024 parameters. Our network model substantially reduces the number of parameters required, by only directly fitting pairwise interactions and getting effective higher order interactions through the local damage term in our model (*f*_*i*_). While a minimal model using our network approach using linear or simple exponential forms for the functions describing rates in our model would have 144 parameters, we have increased this to 188 by using 3rd order power series. This has offered increased flexibility to the model and also improves performance.

Our model generates accurate projections of the average health trajectories of groups of individuals. Taking a group of individuals and simulating them to their deaths, we find the average trajectory generally agrees with the average population data, which shows that model averaged trajectories are quite accurate and are consistent with the assumption that the study population is a random (representative) sample. Note that this does not mean that we can overcome the intrinsic limitations of cross-sectional data, and to validate the accuracy of individual predicted trajectories we would need longitudinal data.

Our model works when separately trained and tested on CSHA and NHANES cross-sectional datasets. This success indicates that our approach should work more broadly with comparable cross-sectional data from other datasets. Nevertheless, we find that performance is somewhat worse on the NHANES dataset. This may be due to the presence of many missing values in the NHANES dataset, the increased age range of the NHANES dataset (20 – 85 years for NHANES vs 65 – 99 for CSHA), or perhaps differing biases in the cohorts studied.

When we predict health trajectories until very old ages (80 - 90 years old), our model tends to estimate slightly higher prevalence than are observed in cross-sectional data (see Figs. 1, S4, and S5), particularly in the NHANES data where baseline measurements are taken further away from the actual age of death than in the CSHA data. One possible explanation for this is that there is a compromise in the model between fitting these trajectories and fitting survival, and the model is attempting to amend this compromise by fitting trajectories well for early ages, then rapidly damaging deficits to induce mortality in individuals to obtain the correct survival predictions. Another possibility is that the damage rates need to rapidly increase from ages 60 to 90, but then slowly taper off of this increase for these older individuals. Fixing this would require more flexible damage rate functions capable of tapering off for very old individuals.

Alternatively, our model could be describing a real acceleration of health decline before death that is not captured in the observational cross-sectional data due the lack of health measurements near death. In other words, these cross-sectional studies could be biased by excluding subjects near death, and our model is correctly including a large increase in the rate of damage near death. Indeed, in longitudinal studies a rapidly rising FI has been shown to identify individuals with a high risk of death within 1 year^30^ – this is called “terminal decline”.^31^ Using such longitudinal data (see below) would allow us to better predict and to better test generated health trajectories for specific individuals, including health near death.

Individual survival is assessed with the C-index.^26^ The C-index evaluates the model’s ability to predict the relative risk of death for pairs of individuals. A C-index of 1 represents perfect predictions, but in practice intrinsic variability of individual mortality will limit the C-index below that in an age and health-dependent way. The availability of individual data will further limit the C-index below that – but generally above 0.5, for an uninformed random guess. We observe C-index values of around 0.6 when stratifying by age, which means that just by using 10 binary deficits, we can predict which of two individuals of the same age lives longer with 60% accuracy. Our model achieves better results than a simple Cox-proportional hazards model^28^ using the Frailty Index, but a larger number of health deficits and more data would further improve our model’s predictions.

Our previous generic network model (GNM) captured population level aging behaviour like Gompertz’ law of mortality and the average increase in the Frailty Index (health decline) vs age,^17, 18^ however the nodes did not correspond to any particular health attribute (i.e. they were generic). Adding more complexity to damage/mortality rates with more flexible functional forms, node-dependent fitting parameters, and a weighted interaction network, we have here been able to represent individual health attributes from observational aging data with specific nodes in our WNM. This has allowed us to model individual health trajectories, including individual survival.

The choice of which deficits to use with our model is arbitrary, the only assumptions we require are that they are binary and not reversible. We do not need the deficits to have strong correlations between them or be good predictors of mortality to capture the sample trends in health trajectories (Figure 1 and Figure 2) or overall survival function (Figure 5A), since these do not show individual predictions but instead captures how well the model overall captures the trends seen in the data. However, the quality of individual predictions for left-out deficits and survival (Figure 4 and Figure 5B) do depend on the deficits chosen, because these are predictions for specific individuals. For independent deficits that are absolutely uninformative of mortality, we would expect AUCs of 0.5 for left-out deficit prediction and a C-index of 0.5 for the mortality prediction.

The only health variables we have included are health deficits that accumulate through damage. Other static or non damage-accumulation variables are often considered in aging studies as well, such as sex,^32–34^ and environmental variables like socioeconomic status,^35^ or lifestyle. Variables that don’t change by damage accumulation but still interact with health deficits can be easily added to the model as network nodes with static values. In this way they could naturally interact with the damage-accumulation health attributes. Similarly, individual non-damage variables that could be deliberately modified – such as physical activity levels^36, 37^ – could be added as nodes with explicit time-dependent values that depend on the individual.

There has been significant work inferring biological networks, using a variety of approaches^38–41^ and at different scales.^42^ In the context of human frailty, previous work has created a network representation of health attributes with measures of association or correlation.^18, 43, 44^ Different methods result in different networks, and thus it has not been clear what underlying association between the deficits a given network represents. Equivalently, it has not been clear how to test a given network representation. In the Supplemental section on Parameter Robustness, we explore the “robustness” of both our network parameters and predictions by sampling an ensemble of parameters around the maximum likelihood estimate.^45^ Supplemental Figure S1 shows that significant deviations from the maximum likelihood parameters still leads to relatively accurate fits of the data – i.e. we obtain robust predictions. However, when we optimized the model several times with different random seeds we show in Figure S2 that we find significantly different network parameterizations each time. Figure S13D shows that even the sign of individual connections is not robust. We conclude that while the model behavior is robust, the network structures themselves are not robustly predicted by the available data.

This lack of robustness of the network is not surprising. Due to the complex interactions between many parameters in our model, we expect that many network parameters are “sloppy”.^46^ This would lead to robust collective behavior of the system with many different combinations of parameters – i.e. many different networks that are consistent with the observed data. Nevertheless, we show in Supplemental Figure S3 that how damage propagates from node to node of the network does have some degree of robustness. We show average damage rates 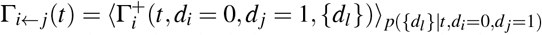 of the *i*th node conditional on prior damage of the *j*th node. This robustness shows that the behavior of our weighted network model (WNM) is robust despite some sloppiness of individual parameters. Indeed, this robust behavior seems necessary to perform well in predicting deficit prevalence and mortality.

Our network model imposes casual mechanisms within the simulation – it assumes that there is a direction to the network weights, and attempts to infer those weights within the assumptions of the model. Since we do not have adequate data to be able to infer true casual relations, these directed links are simply chosen for accurate prediction. A directed link in our model is just defined in terms of prediction: a particular directed connection between two variables is included if it improves prediction accuracy. Similarly, our network connections are not simply correlations between the variables (for those, see Figure S13), but are chosen to improve prediction.

In recent models of disease progression or aging either the mortality is not considered,^15^ or the models require longitudinal data,^47^ or both.^48–50^ Structurally, our model differs from others by using an explicit network describing pairwise interactions, and it uses this network to generate stochastic changes to their health state as they age until death – rather than capturing the dynamics with unobserved latent variables that are harder to interpret. Using discrete health states within our model allows us to simply compare with observed health states using maximum likelihood methods, and allows our success while using only cross-sectional data. That said, our approach could be extended to use longitudinal data for training – and we would expect this to further improve model behavior.

The interpretability of our model structure makes it straightforward to adapt our model to new applications. We can easily generate synthetic tracked health trajectories, or forecast the future trajectories of individuals from specified health states. This means that our model can generate many different stochastic realizations for the same individual after baseline, and can show how differences in possible health trajectories lead to differences in mortality. Another application of our computational approach that would be facilitated by our model structure is to manipulate model individuals to perform “health interventions” on specific observed nodes or sets of nodes. We could then observe the affect of general interventions on health trajectories and mortality. These predictions could then be tested with longitudinal data. This is left for future work.

## Methods

### Model structure

In previous work, we developed a generic network model (GNM)^17, 18^ to study how damage propagation in a network can lead to similar behaviour as observed in aging, in terms of population health and mortality. In this work we expand upon and generalize the GNM to be able to fit the model to individuals with specific observed deficits with a maximum likelihood approach. This allows us to generate synthetic individuals from the model, which age with similar properties as the cohort used to fit the model.

We consider a network of *N* nodes representing binary health attributes. Each node *i* = 1, …, *N* can be in state *d*_*i*_ = 0 for undamaged (healthy) or *d*_*i*_ = 1 damaged (deficit). Nodes in the network undergo stochastic damage transitions (0 → 1) as an individual ages. These transitions occur with rates that depend on the local damage of neighbouring nodes. We call this local damage the “local frailty”, *f*_*i*_.

In the GNM, we measured the local damage around a specific node as the proportion of damaged neighbours, 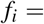 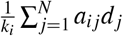, where *a*_*ij*_ is the binary-valued adjacency matrix of an undirected network and *k*_*i*_ = ∑_*j*_ *a*_*ij*_ is the node degree. Damage transitions (0 → 1) between states occurred with rates that depend exponentially on the proportion of damaged neighbours, described by a function 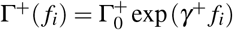 with tunable parameters *γ*^+^ and 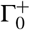. Here we use superscript “+” to denote that this is a damage rate, and correspondingly use “−” to denote repair rates. The baseline rate 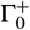 controls the damage rate when *f*_*i*_ = 0, and *γ*^+^ controls how strongly the rate increases with increasing *f*_*i*_. Similar repair transitions were also included (with separate parameters *γ*^−^ and 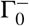), but were found to be negligible. The parameters *γ*^+^ and 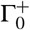 were identical for each node and chosen to fit population mortality rates (Gompertz’ law) and overall health decline (average Frailty Index). For the GNM studies we used *N* = 10^4^ nodes.^17, 18^

In this work, we generalize the GNM to allow the model to represent specific health attributes measured in observed health data as nodes in the network. We generalize the original binary and undirected network to a weighted and directed network, described by a continuous-valued adjacency matrix of weights, *w*_*ij*_. These weights represent the strength of connections between pairs of nodes. We call this a weighted network model (WNM). We use far fewer than 10^4^ nodes in this WNM network, but account for the contribution of these missing nodes by introducing a time-dependent function *μ*_*i*_(*t*) to the local damage of each node, *f*_*i*_. This function *μ*_*i*_(*t*) represents the average contribution to the local damage by the dynamics of the unobserved nodes. This average local damage contribution *μ*_*i*_(*t*) for each node *i* is implemented as a power series in terms of *t* with coefficients 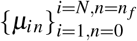, where *n*_*f*_ is a hyperparameter for the number of terms used in the power series. This means our new measure of local damage for the *i*th node is a weighted sum over all of the nodes of the network, with the additional contribution from the *μ*_*i*_(*t*) term,

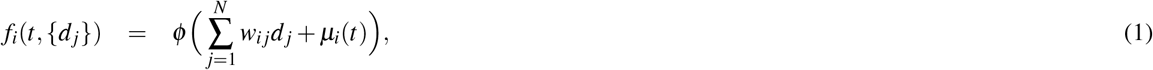

where

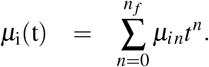

Powers in power series are indexed by *n*, individual deficits or rates are indexed by *i*, and sums over nodes in the network are indexed by *j*. We use this convention of indexing throughout the methods. The function *ϕ* (*x*) = max(*x*, 0) is a “rectifier” or “hinge” function^51^ that clips negative values to zero, resulting in a continuous non-negative function. This can allow strong non-linear behaviour by allowing the function to be able to effectively “turn on” at older ages. The network weights 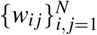 and power series coefficients 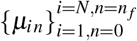 are included as fitting parameters of the model. The coefficients {*μ*_*in*_} are constrained so that *μ*_*i*_(*t*) increases monotonically with age, details are in the Supplemental Information (SI).

The exponential damage rates of the GNM have been replaced by more general power-series in terms of *f*_*i*_, with node-dependent coefficients to allow each node to have a different damage rate. This way, specific nodes in the network are able to represent the distinct behaviour for specific deficits in the observed data (in contrast to the generic network model). This more general damage rate for node *i* is given by,

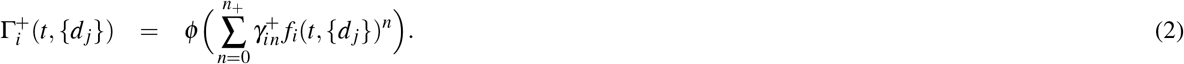

This function describes the damage transition rate from 0 to 1 for node *i*. The power series coefficients 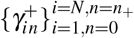 are fitting parameters of the model and *n*_+_ is a hyperparameter that determines the highest-order in the power series. The coefficients 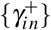 are constrained so that the rate increases monotonically with *f*_*i*_.

Mortality occurs as a separate process with a rate of death that follows the same form with a power series,

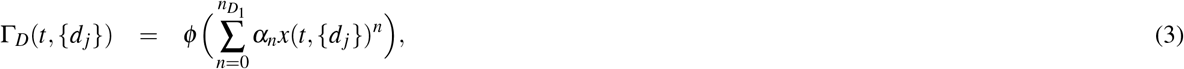

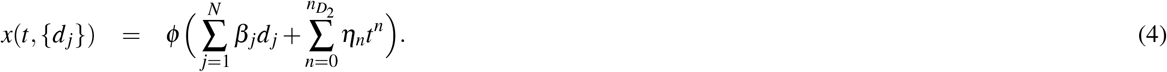

This mortality rate is equivalent to having a single node that corresponds to mortality, and death occurs when it damages (in contrast to the two nodes that were used in the GNM^16, 17^). The measure of local damage *x* that controls mortality is analogous to the local damage *f*_*i*_ for damage rates, and depends on each deficit linearly in a weighted sum. Additionally, it includes an age-dependent deficit-independent function represented as a power series (analogous to *μ*_*i*_(*t*)). This mortality rate uses fitting parameters 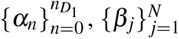, and 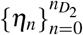 as well as 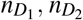 as hyperparameters determining the number of terms in the power series. The coefficients in both Γ_*D*_ and *x* are constrained so that they increase monotonically in *x* and *t*, respectively.

In total the model has 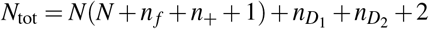 fitting parameters. We restrict parameter values to ensure that *f*_*i*_, 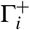, Γ_*D*_, and *x* are all monotonically increasing functions of age. Details of the requisite parameter bounds are in the Supplemental Information (SI). Despite the large number of parameters, we have many more individual observations. We also carefully test predictions for a test population that has small overlap of observed states with our training population (see Supplemental Figure S9). We find no evidence of overfitting.

The model is stochastically simulated by assuming the transition rates describe exponentially distributed waiting times between transitions, and then using an exact event-driven stochastic simulation algorithm (SSA/Kinetic Monte Carlo).^52^ Details of the stochastic simulation are in the SI. For one run of the model until death, i.e. for each synthetic individual, the model outputs death age *t*_*D*_ and all node trajectories, 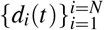 for all *t* between the initial age and mortality. Fully synthetic individuals are started at *t* = 0 with all *d*_*i*_(*t* = 0) = 0, while predicted trajectories for observed individuals are initialized at some *t*_0_ with the completely observed health state at *t*_0_. Due to the exact nature of the SSA, all transition times are precisely resolved in our model data.

### Likelihood

We calculate our likelihood using cross-sectional data. For the *m*th of *M* individuals, we have measurements of health attributes {*d*_*i*_}^(*m*)^ at age *t*^(*m*)^. Instead of death age, we have an observed survival age *a*^(*m*)^ due to right censoring. This is the oldest age that an individual is known to be alive, which can be written 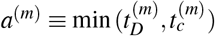, where 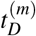 is actual death age and 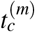 is the censoring age i.e., the age of the individual when they are known to be still alive due to observed health state(s) but after which their mortality is not recorded. We indicate censoring with a binary variable *c*^(*m*)^ = 1, and uncensored with *c*^(*m*)^ = 0. In summary, we consider observed cross-sectional data of the form 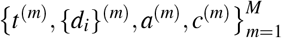.

By simulating synthetic individuals from the model, we sample and estimate the probability 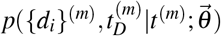 for each individual *m* in the data. We denote all parameters by the vector 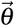. For simplicity we split this probability into two separate parts, representing mortality and health respectively:

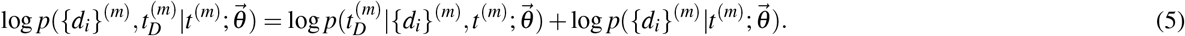

For uncensored individuals, we can calculate their likelihood by using their known death age using Equation 5. For censored individuals, we also need to integrate the mortality term over all possible death ages above the censoring age,

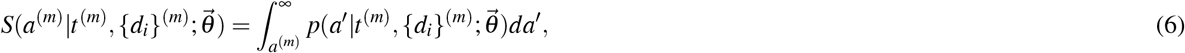

which is the probability of surviving to at least age *a*. Then we can calculate the full log-likelihood,

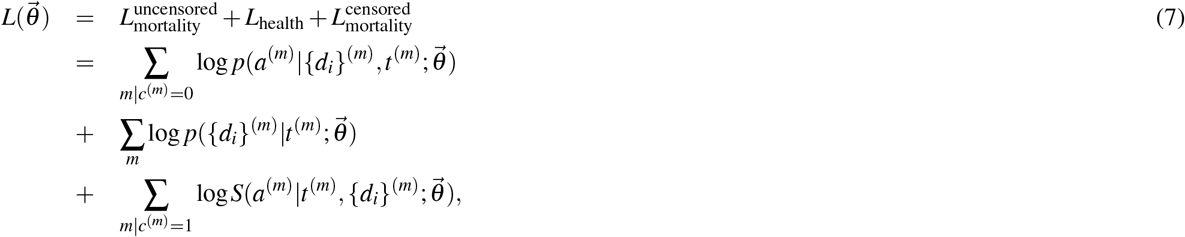

where the last term is added for censored individuals.

For an individual with missing data that does not have the full *N* health attributes measured, we marginalize over the missing values implicitly by sampling all possible combinations of the missing (binary) values. This is done using a synthetic population that has been initialized at *t* = 0 with no damage. Additional details of the likelihood estimation from simulations are in the SI.

### Observed data

We use data from the Canadian Study of Health and Aging (CSHA)^21^ to develop and test our model. The CSHA study used stratified sampling to be a representative sample of the older Canadian population. We use the first wave of the sample with 8547 individuals that range from ages 65 − 99 and death ages that are available within a 6 year censoring window. The mean age is 76 years with a standard deviation of 7 years, the individuals are 60% female, and 78% of individuals have a censored death age. The 10 binary deficits used in the main plots are “Walking difficulty”, “Showering difficulty”, “Phone difficulty”, “Going out difficulty”, “Shopping difficulty”, “Preparing meal difficulty”, “House work difficulty”, “Take medicine difficulty”, “Managing money difficulty”, and “Issues prevent normal activity”. These were chosen by selecting deficits that had large hazard ratios in a Cox proportional hazards analysis,^28^ although any alternate sets of deficits can work, and an alternative set are shown in the SI.

We split the data into a training set of 1020 individuals and a test set of 7527 individuals. We do this such that dividing the training set into 5 year age bins has an approximately uniform age distribution, and the remaining individuals are put into the test set. This balances the training set and ensures no age is “prioritized” in the model training by having a much larger number of individuals.

We validate our conclusions on the National Health And Nutrition Examination Survey (NHANES).^22^ The NHANES dataset used stratified sampling to be a representative sample of the US population. We use a combined sample from the 2003-2004 and 2005-2006 cohorts. The sample has 9504 individuals that range from ages 20 − 85 with death ages that are available within a 10 year censoring window. The mean age is 51 years, with a standard deviation of 20 years, the individuals are 52% female, and 88% of individuals have a censored death age. In the same way as the CSHA data, this data is split into 2352 training individuals and 7151 test individuals.

### Parameter optimization

For each data-set (and choice of health attributes), we maximize the log-likelihood in Equation 7 using particle swarm optimization^53^ in order to train the model and estimate the parameters 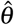. Details of the parameter optimization procedure are in the SI. We use parameter bounds shown in the SI to impose monotonic dependence of damage rates on existing damage. We regularize the fitting as detailed in the SI. We choose hyperparameters *n*_+_ = 4 and 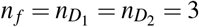. These are the number of terms in our power-series expansions used in damage and mortality functions. The hyperparameters are hand chosen for simplicity. These hyperparameters result in a model with *N*(*N* + 8) + 8 parameters, where N is the number of binary health attributed modelled for each individual. Due to computational demands, this practically limits the size of *N* – here we take *N* = 10 and so have 188 parameters.

## Data availability

NHANES data is available online at https://wwwn.cdc.gov/nchs/nhanes/default.aspx. The 2003-2004 and 2005-2006 cohorts were used for this study. CSHA data is available upon request http://csha.ca/contact_us.asp. The CSHA-1 cohort was used for this study.

## Acknowledgements

We thank ACENET and Compute Canada for computational resources. ADR thanks the Natural Sciences and Engineering Research Council (NSERC) for an operating Grant (RGPIN 2019-05888). KR has operational funding from the Canadian Institutes of Health Research (PJT-156114) and personal support form the Dalhousie Medical Research Foundation as the Kathryn Allen Weldon Professor of Alzheimer Research.

## Author contributions statement

S.F. and A.R. developed the model, S.F. implemented the model and performed all analysis, all authors participated in writing the manuscript. All authors reviewed the manuscript.

## References

1. Kirkwood, T. B. L. Understanding the odd science of aging. Cell 120, 437 – 447 (2005).

2. López-Otín, C., Blasco, M. A., Partridge, L., Serrano, M. & Kroemer, G. The hallmarks of aging. Cell 153, 1194–1217 (2013).

3. Herndon, L. A. et al. Stochastic and genetic factors influence tissue-specific decline in ageing *C. elegans*. Nature 419, 808 – 814 (2002).

4. Kirkwood, T. B. L. & Finch, C. E. The old worm turns more slowly. Nature 419, 794–795 (2002).

5. Mitnitski, A. B., Mogilner, A. J. & Rockwood, K. Accumulation of deficits as a proxy measure of aging. The Sci. World 1, 323–36 (2001).

6. Fried, L. P. et al. Frailty in older adults: Evidence for a phenotype. The Journals Gerontol. Ser. A 56, M146–M157 (2001).

7. Kulminski, A. M., Ukraintseva, S. V., Akushevich, I. V., Arbeev, K. G. & Yashin, A. I. Cumulative index of health deficiencies as a characteristic of long life. J. Am. Geriatr. Soc. 55, 935–940 (2007).

8. Searle, S. D., Mitnitski, A., Gahbauer, E. A., Gill, T. M. & Rockwood, K. A standard procedure for creating a frailty index. BMC Geriatr. 8, 24 (2008).

9. Stubbings, G., Farrell, S., Mitnitski, A., Rockwood, K. & Rutenberg, A. Informative frailty indices from binarized biomarkers. Biogerontology 70, 1–11 (2020).

10. Mitnitski, A. B., Rutenberg, A. D., Farrell, S. & Rockwood, K. Aging, frailty and complex networks. Biogerontology 132, 1–14 (2017).

11. Clegg, A., Young, J., Iliffe, S., Rikkert, M. O. & Rockwood, K. Frailty in elderly people. The Lancet 381, 752–762 (2013).

12. Li, X. et al. Longitudinal trajectories, correlations and mortality associations of nine biological ages across 20-years follow-up. eLife 9, 132 (2020).

13. Mitnitski, A. B., Mogilner, A. J., MacKnight, C. & Rockwood, K. The mortality rate as a function of accumulated deficits in a frailty index. Mech. Ageing Dev. 123, 1457–1460 (2002).

14. Rockwood, K. & Mitnitski, A. Frailty in relation to the accumulation of deficits. J. Gerontol. Med. Sci. 62A, 722–727 (2007).

15. Pierson, E., Koh, P. W., Hashimoto, T., Koller, D. & Liang, P. Inferring multidimensional rates of aging from cross-sectional data. Proc Mach Learn. Res 89, 97–107 (2019).

16. Taneja, S., Mitnitski, A. B., Rockwood, K. & Rutenberg, A. D. Dynamical network model for age-related health deficits and mortality. Phys. Rev. E 93, 022309 (2016).

17. Farrell, S. G., Mitnitski, A. B., Rockwood, K. & Rutenberg, A. D. Network model of human aging: frailty limits and information measures. Phys. Rev. E 94, 052409 (2016).

18. Farrell, S. G., Mitnitski, A. B., Rockwood, K. & Rutenberg, A. D. Probing the network structure of health deficits in human aging. Phys. Rev. E 98, 032302 (2018).

19. Kirkwood, T. B. L. Deciphering death: a commentary on gompertz (1825) ‘on the nature of the function expressive of the law of human mortality, and on a new mode of determining the value of life contingencies’. Philos. Transactions Royal Soc. B 370, 20140379 (2015).

20. Rutenberg, A. D., Mitnitski, A. B., Farrell, S. G. & Rockwood, K. Unifying aging and frailty through complex dynamical networks. Exp. Gerontol. 107, 126–129 (2018).

21. Canadian Study of Health and Aging Working Group. Canadian study of health and aging: study methods and prevalence of dementia. Can. Med. Assoc. J. 150, 899 (1994).

22. Centers for Disease Control and Prevention National Center for Health Statistics. National health and nutrition examination survey data (Updated 2014).

23. Bennett, S., Song, X., Mitnitski, A. & Rockwood, K. A limit to frailty in very old, community-dwelling people: a secondary analysis of the Chinese longitudinal health and longevity study. Age Ageing 42, 372–377 (2013).

24. Armstrong, J. J., Mitnitski, A., Launer, L. J., White, L. R. & Rockwood, K. Frailty in the Honolulu-Asia Aging Study: deficit accumulation in a male cohort followed to 90% mortality. Journals Gerontol. Ser. A: Biol. Sci. Med. Sci. 70, 125–131 (2015).

25. Kaplan, E. L. & Meier, P. Nonparametric estimation from incomplete observations. J. Am. Stat. Assoc. 53, 457–481 (1958).

26. Harrell Jr, F. E., Califf, R. M., Pryor, D. B., Lee, K. L. & Rosati, R. A. Evaluating the yield of medical tests. The J. Am. Med. Assoc. 247, 2543–2546 (1982).

27. Antolini, L., Boracchi, P. & Biganzoli, E. A time-dependent discrimination index for survival data. Stat. Medicine 24, 3927 – 3944 (2005).

28. Cox, D. R. Regression models and life-tables. J. Royal Stat. Soc. Ser. B 34, 187–200 (1972).

29. Graf, E., Schmoor, C., Sauerbrei, W. & Schumacher, M. Assessment and comparison of prognostic classification schemes for survival data. Stat. Medicine 18, 2529–2545 (1999).

30. Stow, D., Matthews, F. E. & Hanratty, B. Frailty trajectories to identify end of life: a longitudinal population-based study. BMC Medicine 16, 1–7 (2018).

31. Cohen-Mansfield, J., Skornick-Bouchbinder, M. & Brill, S. Trajectories of end of life: A systematic review. Journals Gerontol. - Ser. B Psychol. Sci. Soc. Sci. 73, 564–572 (2018).

32. Puts, M. T. E., Lips, P. & Deeg, D. J. H. Sex differences in the risk of frailty for mortality independent of disability and chronic diseases. J. Am. Geriatr. Soc. 53, 40–47 (2005).

33. Mitnitski, A. et al. Relative fitness and frailty of elderly men and women in developed countries and their relationship with mortality. J. Am. Geriatr. Soc. 53, 2184–2189 (2005).

34. Gordon, E. H., Peel, N. M., Theou, O., Howlett, S. E. & Hubbard, R. E. Sex differences in frailty: A systematic review and meta-analysis. Exp. Gerontol. 89, 30–40 (2017).

35. Andrew, M. K., Mitnitski, A. B. & Rockwood, K. Social vulnerability, frailty and mortality in elderly people. PLOS One 3, e2232 (2008).

36. Fried, L. P. Interventions for human frailty: Physical activity as a model. Cold Spring Harb. Perspectives Medicine 6, a025916 (2016).

37. Rogers, N. T. et al. Physical activity and trajectories of frailty among older adults: Evidence from the english longitudinal study of ageing. PLOS One 12, e0170878 (2017).

38. Butte, A. J. & Kohane, I. S. Mutual information relevance networks: functional genomic clustering using pairwise entropy measurements. Pac. Symp. on Biocomput. 5, 415–426 (2000).

39. Margolin, A. A. et al. ARACNE: An algorithm for the reconstruction of gene regulatory networks in a mammalian cellular context. BMC Bioinforma. 7, S7 (2006).

40. Zhang, B. & Horvath, S. A general framework for weighted gene co-expression network analysis. Stat. Appl. Genet. Mol. Biol. 4, 1 (2005).

41. Shen, Z., Wang, W.-X., Fan, Y., Di, Z. & Lai, Y.-C. Reconstructing propagation networks with natural diversity and identifying hidden sources. Nat. Commun. 5, 4323 (2014).

42. Natale, J. L., Hofmann, D., Damian G, H. & Nemenman, I. Reverse-engineering biological networks from large data sets. Quant. Biol. Theory, Comput. Methods, Model. Chapter 10 (2017).

43. Mitnitski, A. B., Graham, J. E., Mogilner, A. J. & Rockwood, K. Frailty, fitness and late-life mortality in relation to chronological and biological age. BMC Geriatr. 2, 1 (2002).

44. García-Peña, C. et al. Network analysis of frailty and aging: Empirical data from the Mexican Health and Aging Study. Exp. gerontology 128, 110747 (2019).

45. Zhou, B., Hofmann, D., Pinkoviezky, I., Sober, S. J. & Nemenman, I. Change, long tails, and inference in a non-Gaussian, Bayesian theory of vocal learning in songbirds. Proc. Natl. Acad. Sci. 115, E8358 (2018).

46. Gutenkunst, R. N. et al. Universally sloppy parameter sensitivities in systems biology models. PLoS Comput. Biol 3, 10 (2007).

47. Lim, B. & van der Schaar, M. Disease-atlas: Navigating disease trajectories using deep learning. Proceeding Mach. Learn. Res. 85, 137–160 (2018).

48. Schulam, P. & Saria, S. A framework for individualizing predictions of disease trajectories by exploiting multi-resolution structure. In Advances in Neural Information Processing Systems, 748–756 (Johns Hopkins University, Baltimore, United States, 2015).

49. Alaa, A. M. & van der Schaar, M. Forecasting individualized disease trajectories using interpretable deep learning. arXiv arXiv:1810.10489v1 (2018).

50. Fisher, C. K., Smith, A. M. & Walsh, J. R. Machine learning for comprehensive forecasting of Alzheimer’s Disease progression. Sci. Reports 9, 1–14 (2019).

51. Breiman, L. Hinging hyperplanes for regression, classification, and function approximation. Biogerontology 39, 999–1013 (1993).

52. Gillespie, D. T. Exact stochastic simulation of coupled chemical reactions. The J. Phys. Chem. 81, 25 (1977).

53. Kennedy, J. & Eberheart, R. Particle swarm optimization. Proc. IEEE Int. Conf. on Neural Networks IV, 1942–1948 (1995).

