## Supplemental information for "Generating synthetic aging trajectories with a weighted network model using cross-sectional data"

October 26, 2020

#### S1 Simulation details

##### S1.1 Stochastic simulation

We exactly simulate the model for each individual using a standard stochastic simulation algorithm [1]. Two separate rate processes are simulated for damage and mortality. The first process for damage uses the rate in Eqn. [2] in the main text. (Supplemental equations, figures, and sections are denoted with 'S'. All supplemental references are listed at the end of the supplemental information.) The second process uses the mortality rate in Eqn. [3].

Since our rates are time-dependent we use an exact rejection sampling method, called the "attempt time algorithm" [2]. We set the window for possible events as $T = 5$ years, and re-update this maximum horizon as it is passed. Since we have constrained all rates to be monotonic in time (see below for details), we can easily determine an upper bound on the rates, necessary to implement the attempt time algorithm: $\Gamma^{+}_{\max} = \sum_i (1 - d_i)\Gamma^{+}_i(t + T, \{d_j\})$ bounds the total damage rate of all undamaged nodes, while $\Gamma^{D}_{\max} = \Gamma_D(t + T, \{d_j\})$ bounds the mortality rate.

At each step of the algorithm, the event is chosen from the process with the smallest time-to-event, i.e., damage occurs until the time to death is smaller than the time to next damage event. Within the damage process, we choose which event using a computationally efficient tree-based approach [3]. When the mortality event occurs, the algorithm terminates for that individual.

##### S1.2 Estimating likelihood

To calculate the likelihood in Eqn. [7], we run $\mathbb{S}$ simulations with the same set of parameters (with $\mathbb{S} > 10^6$ individuals). We use a discrete kernel density estimate of the health term of the likelihood,

$$p(\{d_i\}^{(m)}|t^{(m)};\vec{\theta}) = \frac{1}{\mathbb{S}(t^{(m)})} \sum_{s=1}^{\mathbb{S}(t^{(m)})} K(\{d_i\}^{(s)}, \{d_i\}^{(m)}; \lambda^{(m)}), \tag{S1}$$

where $\mathbb{S}(t^{(m)})$ is the number of simulated individuals alive at age $t^{(m)}$. The superscript $(s)$ indicates a simulated individual and $(m)$ an observed individual from the data. We use the Racine and Li kernel [4],

$$K(\{d_i\}^{(s)}, \{d_i\}^{(m)}; \lambda^{(m)}) = \begin{cases} \frac{1}{1+\lambda^{(m)}}, & \sum_i |d_i^{(s)} - d_i^{(m)}| = 0,\ t^{(s)} = t^{(m)} \\ \frac{\lambda^{(m)}}{1+\lambda^{(m)}}, & \sum_i |d_i^{(s)} - d_i^{(m)}| = 1,\ t^{(s)} = t^{(m)} \\ 0, & \text{otherwise.} \end{cases} \tag{S2}$$

This allows individuals that differ in damage by 1 deficit to contribute to the likelihood, as determined by an individual bandwidth $\lambda^{(m)}$.

The bandwidth $\lambda^{(m)}$ is selected by minimizing the mean-squared error [5],

$$\lambda^{(m)} = \left[1 + \frac{\mathbb{S}(t^{(m)})((p^{(m)})^2 + (1-p^{(m)})^2)}{2p^{(m)}(1-p^{(m)})}\right]^{-1}, \tag{S3}$$

where $p^{(m)}$ is the empirical frequency estimate of the distribution (obtained using bandwidth $\lambda = 0$). The bandwidth $\lambda^{(m)}$ is individual-dependent, so that the well sampled individuals will have a small bandwidth, and poorly sampled individuals have a larger bandwidth to reduce noise.

Both the censored and uncensored mortality terms of the likelihood are calculated by binning the death ages of simulated individuals that match the observed data ($\sum_i |d_i^{(s)} - d_i^{(m)}| = 0,\ t^{(s)} = t^{(m)}$) with 1 year bins.

#### S1.3 Regularization

To have an increase in mortality rate with decreasing health, the deficit mortality contributions $\beta_j$ in Eqn. [4] should all be positive. Without such a bound, we have observed that the optimization sometimes converges to positive and negative $\beta_j$ values scattered around zero, which leads to the deficit contribution to the mortality rate being negligible, and gives uniform mortality for all individuals. However in contrast, setting a strict bound $\beta_j \geq 0$ causes the optimization to converge to parameters which perform poorly with respect to the health trajectories. Instead, we have found that a soft penalty behaves well when added to the log-likelihood to penalize negative values,

$$+C\sum_j \min(0, \beta_j). \tag{S4}$$

Values of $\beta_j \sim 1$ give a significant mortality contribution, and log-likelihood values are estimated with a stochastic noise of $\pm 10$, so we choose $C = 100$ by hand to achieve a moderate regularizing effect – and we find that the optimization performs well with this choice.

#### S1.4 Parameter bounds

To ensure Eqns. [1], [2], [3], and [4] are monotonically increasing for all $t \geq 0$, we bound the parameters so that their time-derivatives have only negative real or complex roots with respect to $t$ or $f_i$. For hyperparameters $n_+ = 4$, and $n_f = n_{D_1} = n_{D_2} = 3$, the required bounds are shown in Table S1.

#### S1.5 Parameter optimization

Optimization is done with particle swarm optimization (PSO) [6]. PSO is derivative-free and highly parallelizable. We use the standard version:

$$\begin{aligned} \mathbf{v}_{i,t+1} &= \omega \mathbf{v}_{i,t} + c_p u_p(\vec{\theta}^p_{i,t} - \vec{\theta}_{i,t}) + c_g u_g(\vec{\theta}^g_t - \vec{\theta}_{i,t}) && \text{(S5)} \\ \vec{\theta}_{i,t+1} &= \vec{\theta}_{i,t} + \mathbf{v}_{i,t+1}. && \text{(S6)} \end{aligned}$$

Table S1: Parameter bounds during optimization.

| Equation | Parameter bounds |
|---|---|
| Eqn. [1]: $f_i(t, \{d_i\})$ | $\mu_{i1} \geq 0$, $\mu_{i3} \geq 0$, $\mu_{i2} \geq -\sqrt{3\mu_{i1}\mu_{i3}}$ |
| Eqn. [2]: $\Gamma_i^+(t, \{d_i\})$ | $\gamma_{i1} \geq 0$, $\gamma_{i3} \geq 0$, $\gamma_{i2} \geq -\sqrt{3\gamma_{i1}\gamma_{i3}}$, $\gamma_{i4} \geq 0$ |
| Eqn. [3]: $\Gamma_D(t, \{d_i\})$ | $\alpha_1 \geq 0$, $\alpha_3 \geq 0$, $\alpha_2 \geq -\sqrt{3\alpha_1\alpha_3}$ |
| Eqn. [4]: $x(t, \{d_i\})$ | $\eta_1 \geq 0$, $\eta_3 \geq 0$, $\eta_2 \geq -\sqrt{3\eta_1\eta_3}$ |

The parameter values for the $i$th particle at an iteration $t$ are represented as $\theta_{i,t}$ and the velocity for this particle as $\mathbf{v}_{i,t}$. The current best set of parameters found by particle $i$ is $\theta^p_{i,t}$, and the current global best set of parameters is $\theta^g_t$. We randomly sample $u_p$, $u_g$ uniformly from $[0, 1]$. We set $\omega = 0.7$, $c_p = c_g = 2$ and use $100 - 200$ particles, depending on compute resources available. The optimization typically converges in 100-200 iterations.

Parameter bounds from section S1.4 are implemented by rescaling the velocity components $v_{i,k,t}$ with a hyperbolic method so that parameters never exceed their upper bounds $U_k$ and lower bounds $L_k$ [7],

$$v_{i,k,t+1} = \frac{v_{i,k,t+1}}{1 + \frac{v_{i,k,t+1}}{U_k - \theta_{i,k,t}}}, v_{i,k,t+1} > 0, \tag{S7}$$
$$v_{i,k,t+1} = \frac{v_{i,k,t+1}}{1 - \frac{v_{i,k,t+1}}{\theta_{i,k,t} - L_k}}, v_{i,k,t+1} < 0.$$

Our objective function is stochastic, so to avoid optimizing to the noise, every 10 PSO iterations we recalculate the global maximum likelihood for the global best set of parameters $\theta^g_t$, and the individual particle maximum likelihood values for each particle's best set of parameters $\theta^p_{i,t}$.

#### S2 Parameter robustness

To examine parameter robustness, we select new parameter sets by randomly selecting 25% of parameters to randomly perturb from their maximum likelihood values $\hat{\theta}_d$ within the range $\theta_d \in [0.1\hat{\theta}_d, 10\hat{\theta}_d]$.

Broadly following the approach of [8], we weight these perturbed parameters by calculating an approximate log-likelihood by using only the $\chi^2$ of first and second order joint distributions of health and population survival. This amounts to a Gaussian approximation to the log-likelihood,

$$\begin{aligned}
\mathcal{L}(\theta) &\simeq -\frac{1}{N(T_{\max}-T_{\min})}\sum_{i=1}^{N}\int_{T_{\min}}^{T_{\max}}\Big(\frac{\hat{p}(d_i=1|t;\theta)-p(d_i=1|t)}{\sigma_{\hat{p}}(t)}\Big)^2 dt \qquad \text{(S8)}\\
&- \frac{2}{N(N-1)(T_{\max}-T_{\min})}\sum_{i,j\neq i}\int_{T_{\min}}^{T_{\max}}\Big(\frac{\hat{p}(d_i=1,d_j=1|t;\theta)-p(d_i=1,d_j=1|t)}{\sigma_{\hat{p}}(t)}\Big)^2 dt\\
&- \frac{1}{N(N-1)(T_{\max}-T_{\min})}\sum_{i,j}\int_{T_{\min}}^{T_{\max}}\Big(\frac{\hat{p}(d_i=1,d_j=0|t;\theta)-p(d_i=1,d_j=0|t)}{\sigma_{\hat{p}}(t)}\Big)^2 dt\\
&- \frac{1}{T_{\max}-T_{\min}}\int_{T_{\min}}^{T_{\max}}\Big(\frac{\langle\hat{S}(a|\{d_i\},t;\theta)\rangle-S(a|t)}{\sigma_{\hat{S}}(t)}\Big)^2 dt.
\end{aligned}$$

We use this estimate of $\mathcal{L}$ to efficiently estimate any $f(\theta)$:

$$\langle f(\theta)\rangle = \frac{\sum_{\theta_i} e^{\mathcal{L}(\theta)} f(\theta)}{\sum_{\theta_i} e^{\mathcal{L}(\theta)}}, \qquad \text{(S9)}$$

$$\sigma_f^2 = \langle f(\theta)^2\rangle - \langle f(\theta)\rangle^2. \qquad \text{(S10)}$$

In Fig. S1 we show deficit prevalence and survival averaged over likelihood-weighted parameterizations via Eqns. S9 and S10. These average predictions are quite close to the maximum likelihood estimates, which indicate that prevalences and mortality are robust predictions of our modelling approach.

In contrast, simple measures of the network structure do not appear to be robust. In Fig. S2 we perform the optimization 13 times, and the compare network degrees for the 10 nodes of our network. We find a broad ranges of degrees for each node, with significant overlap of these ranges between nodes. This indicates that though the behavior of the networks is similar (see e.g. Fig. S3), the network structures themselves are not robustly predicted by the available data.

#### Supplemental figures

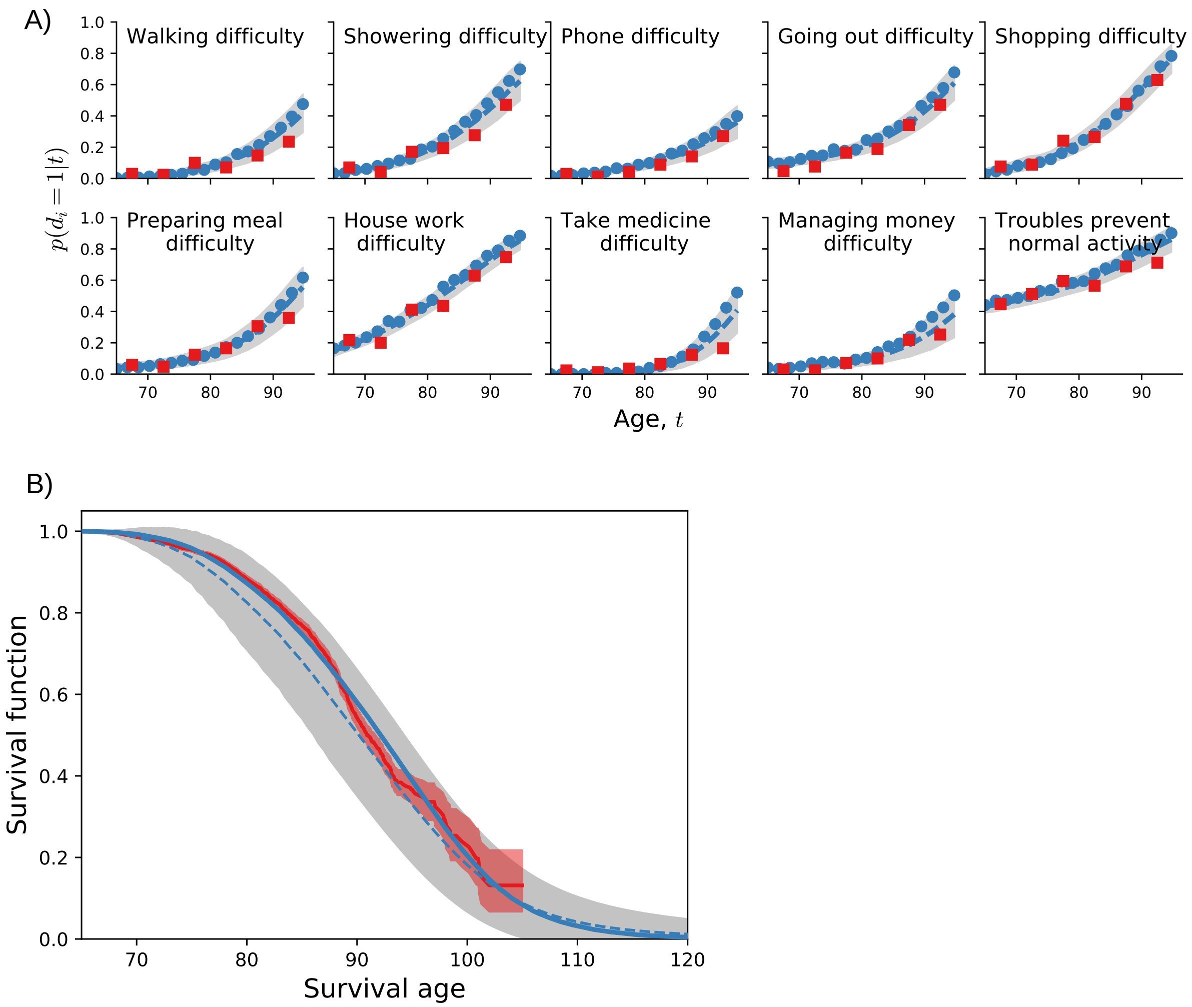


Fig S1: **Prediction robustness.** A) Average predicted trajectories of ten deficit prevalences for individuals from the test data aged $65 - 70$ surviving past 70 (blue points). Observed CSHA prevalence is shown in red squares. B) Model population average survival function (blue line) and CSHA Kaplan-Meier population survival function (red line and shaded region). For both A) and B), the dashed blue lines are averages of the prevalence or individual survival curves over different parameter sets (Eqn. S9), the shaded grey region shows one standard deviation (Eqn. S10) from this average. Agreement of the shaded grey regions with our average results indicates that prevalences and survival are robustly predicted by our approach.

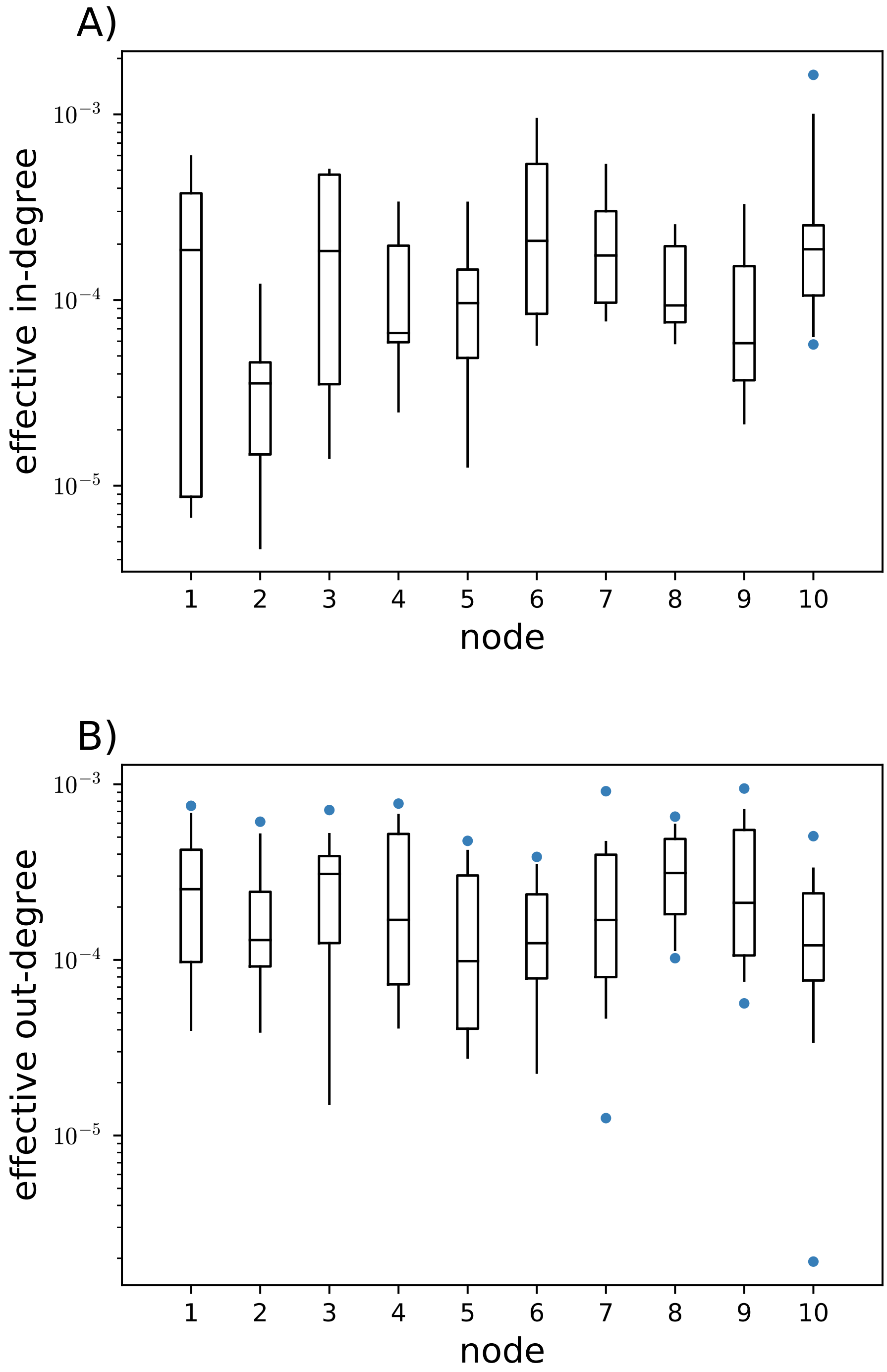


Fig S2: **Network sloppiness.** Maximum likelihood effective network degrees for 13 numerical optimizations starting from different random seeds. Boxes show the interquartile range and whiskers show the 2.5% and 97.5% values. The network structure does not appear to be robust (note the log-scale). A) Effective in-degree $k_i^{\text{in}} = \sum_j w_{ij}\gamma_{i1}^{+}$ for each node. B) Effective out-degree $k_j^{\text{out}} = \sum_i w_{ij}\gamma_{i1}^{+}$ for each node.

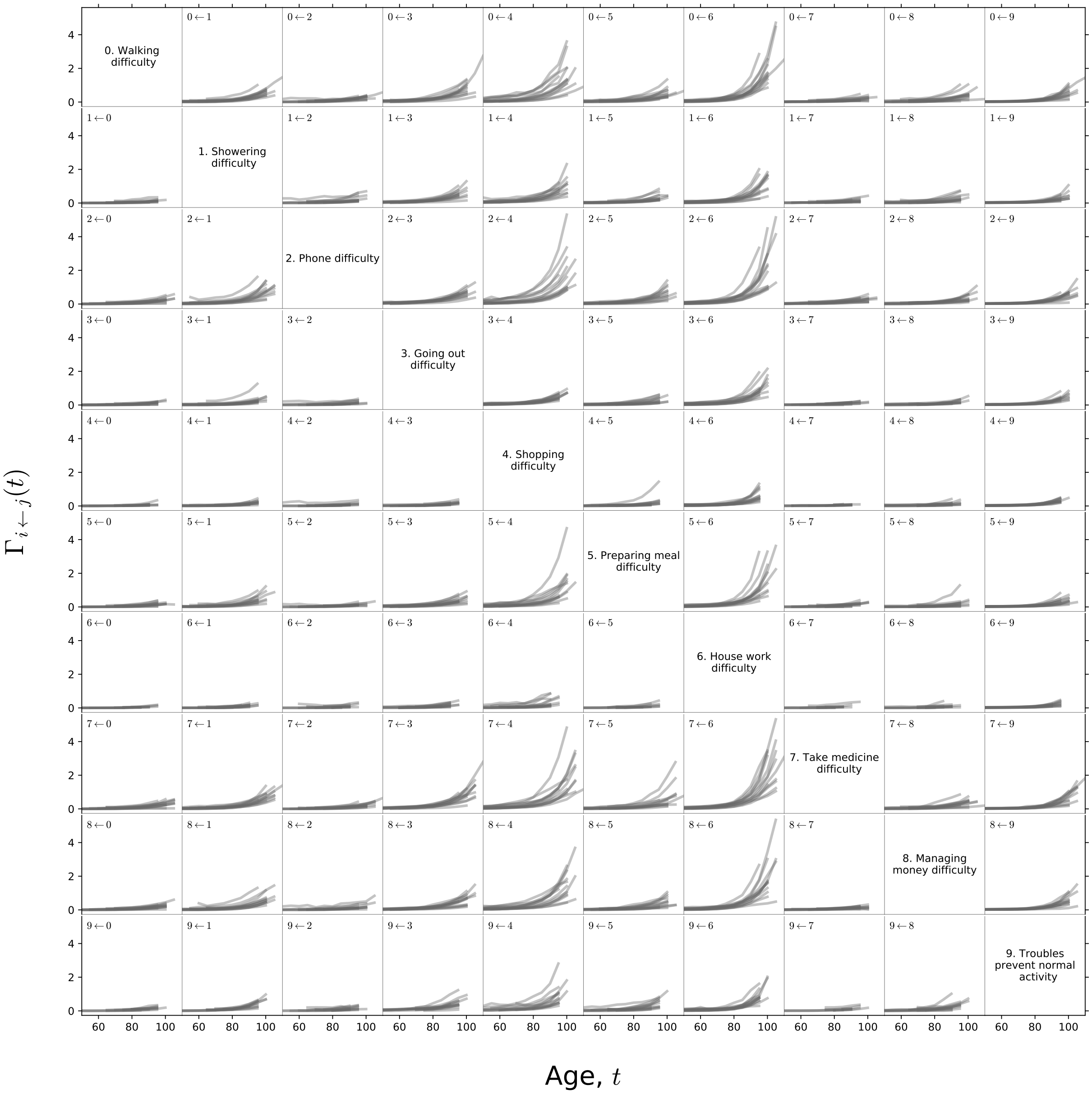


Fig S3: **Damage rate robustness.** Average damage rate $\Gamma_{i\leftarrow j}(t) = \langle \Gamma_i^+(t, d_i = 0, d_j = 1, \{d_l\}) \rangle_{p(\{d_l\}|t, d_i=0, d_j=1)}$ of node $i$ given that node $j$ is damaged, vs age. Each curve is averaged over combinations of the other node states, weighted by their probability of occurring in the simulation. Each intersecting row and column represents the indicated node labelled on the diagonal, and the direction of the links is indicated by labels on each subplot. Each subplot has 13 rate curves, each with different estimated parameters for a different starting seed. The qualitative agreement of these curves for particular pairs of nodes indicates that these detailed damage rates are qualitatively robust (note the linear scale), even though the effective degrees of the nodes are not (see Fig. S2).

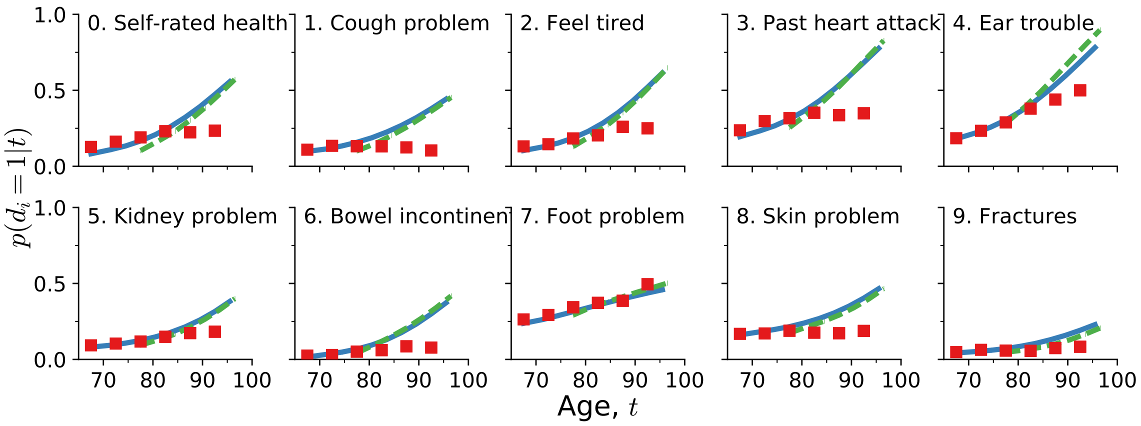


Fig S4: **Alternate deficit predicted trajectories.** Average predicted trajectories of deficit prevalence for individuals from the test data aged $65 - 70$ surviving past 70 (solid blue lines) and aged $75 - 80$ surviving past 80 (dashed green lines). An alternate set of deficits are used as compared to Fig. 1, in the main text. CSHA prevalence is shown in with red squares, standard errors are smaller than the point size.

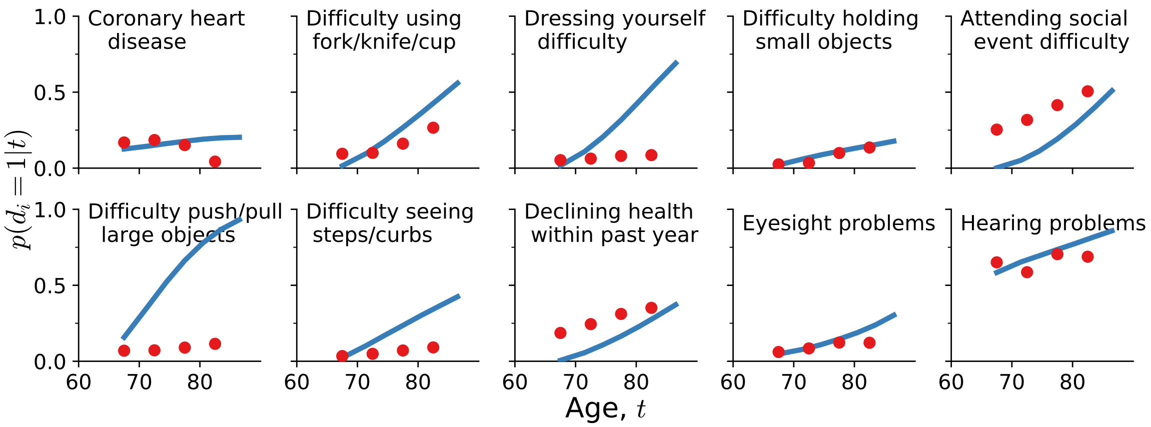


Fig S5: **NHANES predicted trajectories.** Average predicted trajectories of deficit prevalence for individuals from the test data aged $65 - 70$ surviving past 70 (solid blue lines). Observed NHANES prevalence is shown with red circles; standard errors are smaller than the point size. Here predictions do not always start at value from the observational data (comparing the agreement for the first red point with Fig. S4) because there is missing data that we impute by simulating from birth.

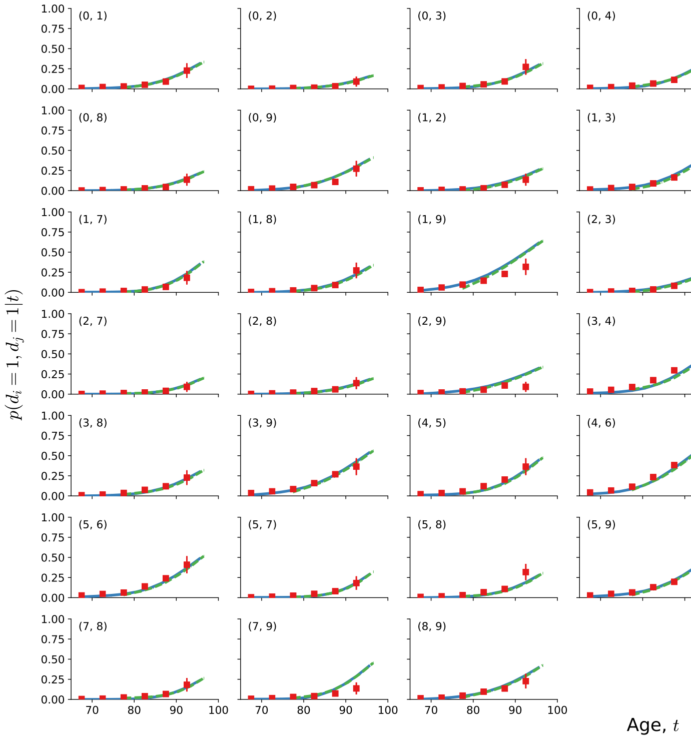


Fig S6: **Full pair trajectories.** Average predicted trajectories of pairwise deficit prevalence for individuals from the test data aged $65-70$ surviving past 70 (solid blue lines) and aged $75-80$ surviving past 80 (dashed green lines). Subplot titles indicate the two deficits included in the pairwise prevalence, with numbers corresponding to titles in Fig. 1, in the main text. All pairs are shown. CSHA data is shown by red squares; errorbars represent standard errors of the pairwise prevalence.

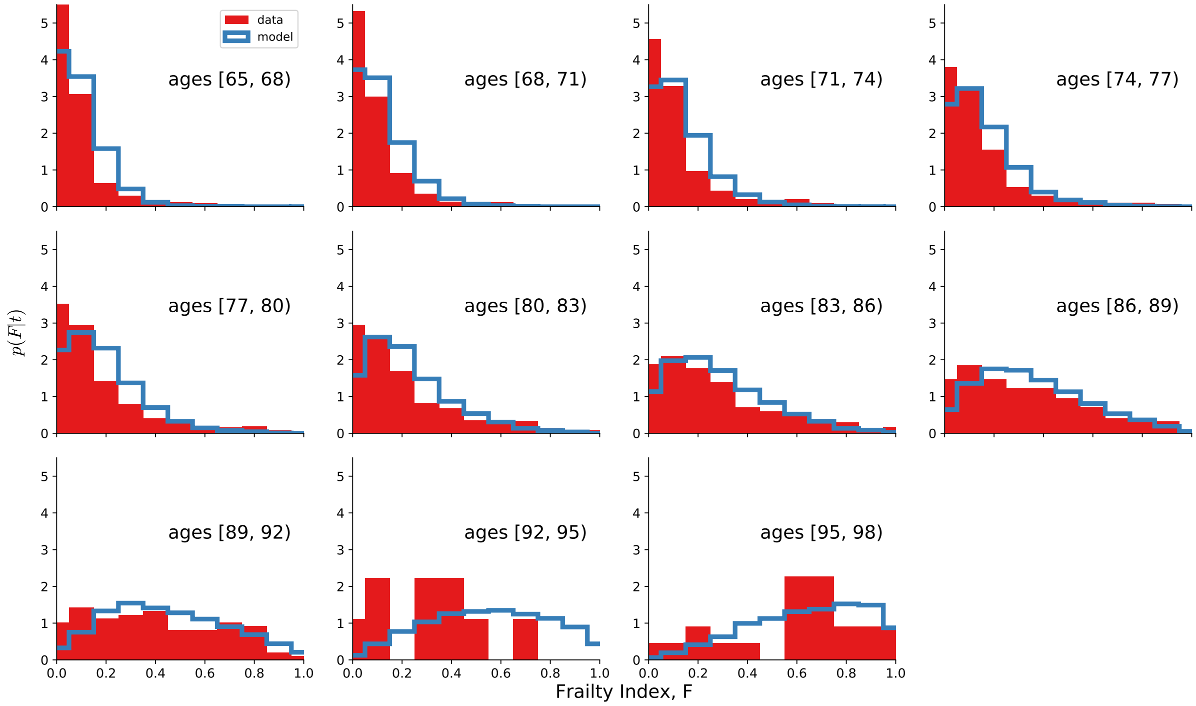


Fig S7: **Population Frailty Index distributions.** Distribution of the Frailty Index, $F = \sum_{i=1}^{N} d_i/N$ for different age ranges for the model (unfilled blue histograms) and the CSHA data (filled red histograms). Note that the maximum FI for small numbers of deficits (in this case $N = 10$) is 1, as is reached in the CSHA data at later ages. Recovering this FI limit with model data, as well as broader consistency with the CSHA distributions, confirms that we capture the heterogeneity of the population as individuals age.

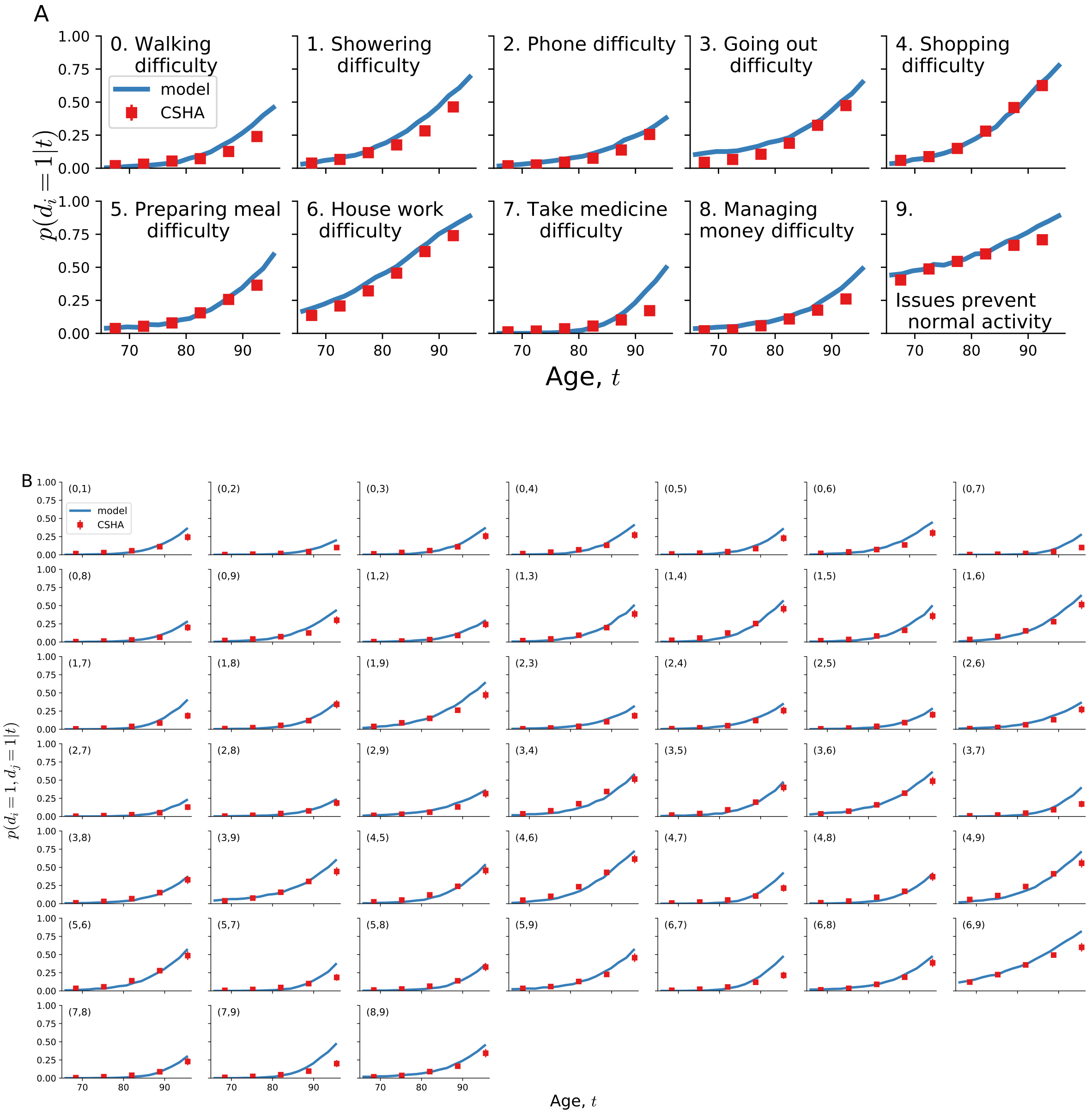


Fig S8: **Simulated population trajectories from birth.** A) Predicted deficit prevalence $\hat{p}(d_i = 1|t)$ vs age $t$ in our model simulated from zero damage at birth (blue lines) and binned prevalence from the CSHA (red squares). B) Predicted pair prevalence $\hat{p}(d_i = 1, d_j = 1|t)$ vs age in our model simulated from zero damage at birth (blue lines) is compared with binned pair prevalence from the CSHA data (red squares). Standard errors of the data are smaller than the point size.

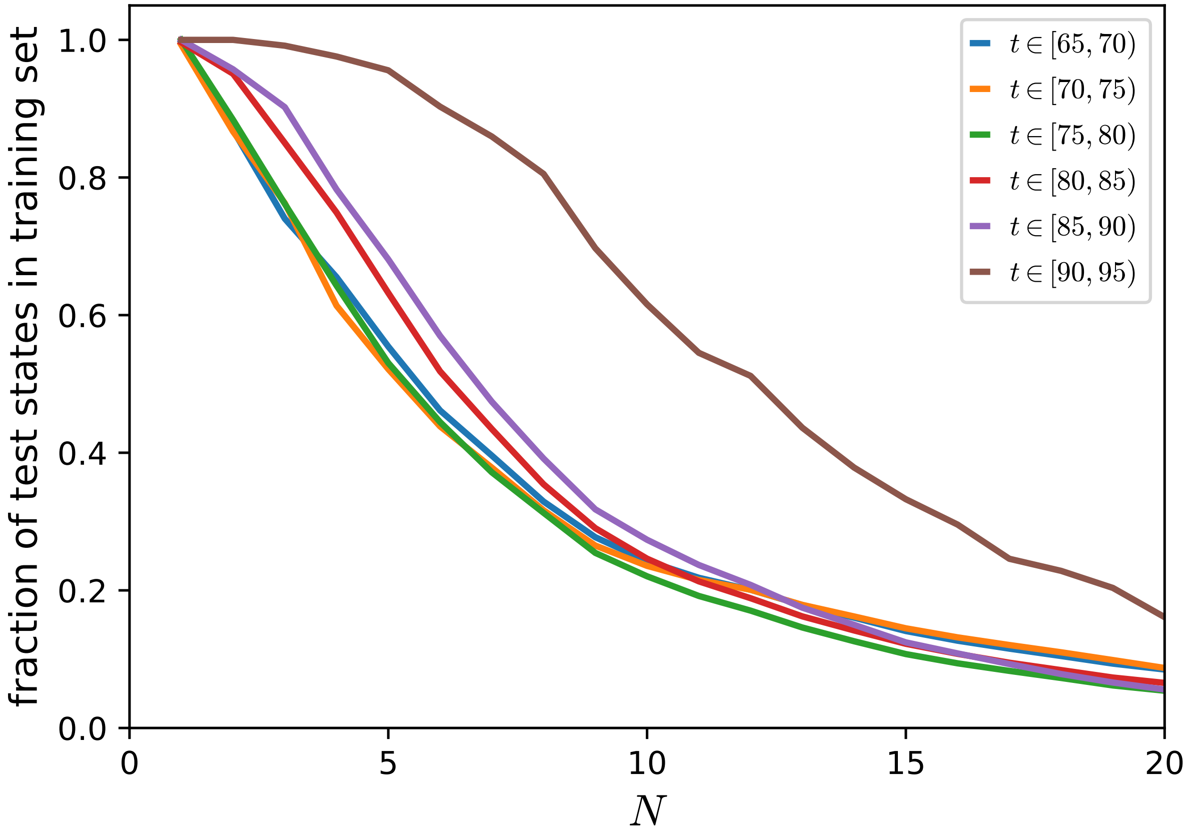


Fig S9: **CSHA state overlap** Fraction of health states $\{d_i\}_{i=1}^N$ in the test set for different ranges of ages (as indicated by the legend) that are also in the training set at any age vs. $N$, the total number of health attributes (nodes) in our model. We average over permutations of the order that the deficits are chosen. We show that a significant fraction of the states in the test set are not in training set for $N = 10$, except for the oldest individuals aged $90 - 95$.

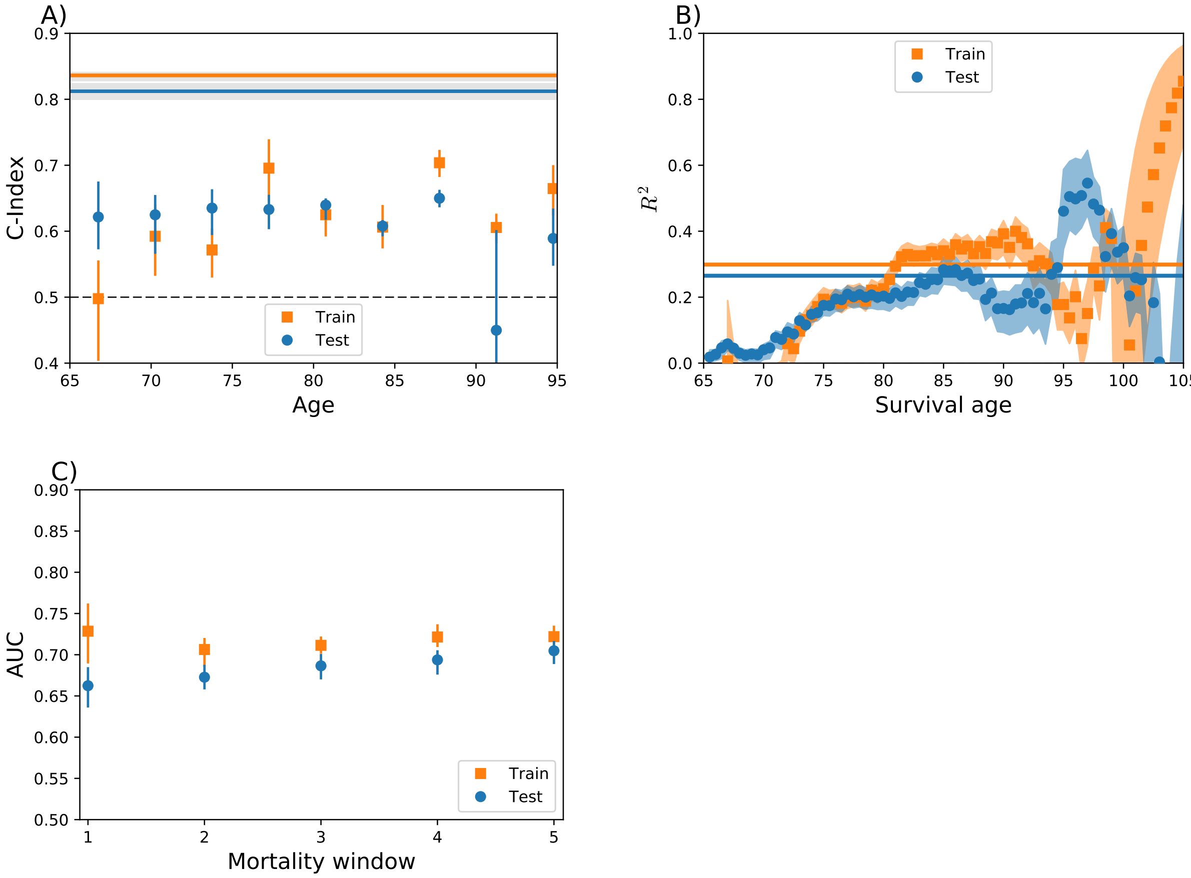


Fig S10: **Training set mortality predictions.** We observe similar performance between train (orange squares and lines) and test (blue circles and lines) CSHA data, indicating a lack of overfitting. All errorbars and shaded regions show 95% confidence intervals. A) Survival C-Index stratified by age (points) and unstratified (solid lines). The baseline for random predictions is at $C = 0.5$ (dashed black line). B) Explained variance of survival $R^2(a) = 1 - \mathrm{BS}(a)/\mathrm{BS}_0(a)$ from the Brier score [9] is shown as points, with uncertainties indicated with corresponding shaded regions. Baseline Brier Score is calculated with the health-independent Kaplan-Meier estimate of the survival function, $S(a)$. Integrated $R^2$ over all ages is shown as solid lines. Values above zero show improvement above baseline. C) ROC AUC for the prediction of mortality within a window of time (in years). The baseline for random predictions is at $AUC = 0.5$.

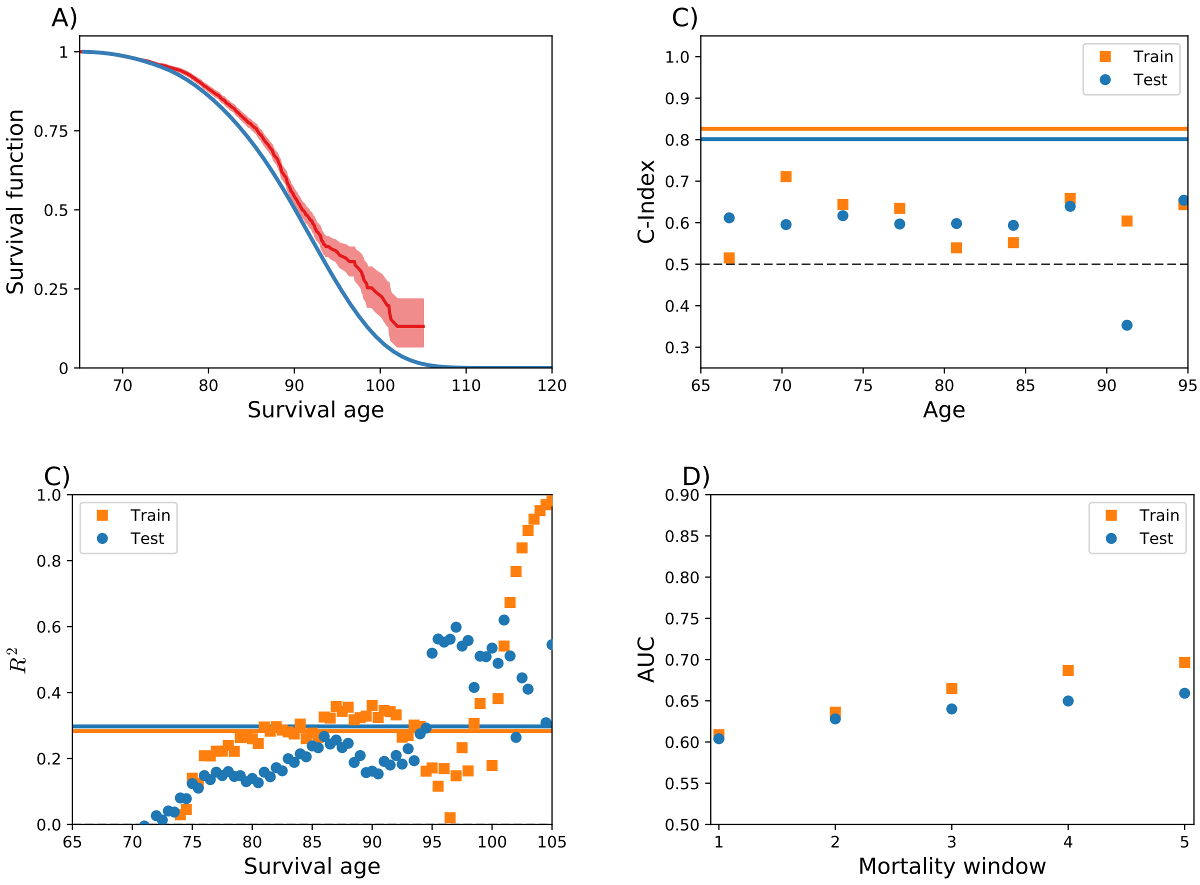


Fig S11: **Alternate deficits mortality predictions.** Survival predictions with alternative deficits used in Supplemental Fig. S4 with CSHA data. Compare with Fig. S10. Model predictions work similarly well with these alternate deficits. A) Model population average survival function (blue) and Kaplan-Meier population survival function (red) with a 95% confidence interval. B) Survival C-Index stratified by age (points) and unstratified (solid lines). The baseline for random predictions is $C = 0.5$ (black dashed line). C) Explained variance $R^2$ from the Brier score. Integrated $R^2$ is shown as solid lines. Higher values greater than zero mean better predicted survival curves. D) ROC AUC for the prediction of mortality within a window of time (in years) for CSHA data.

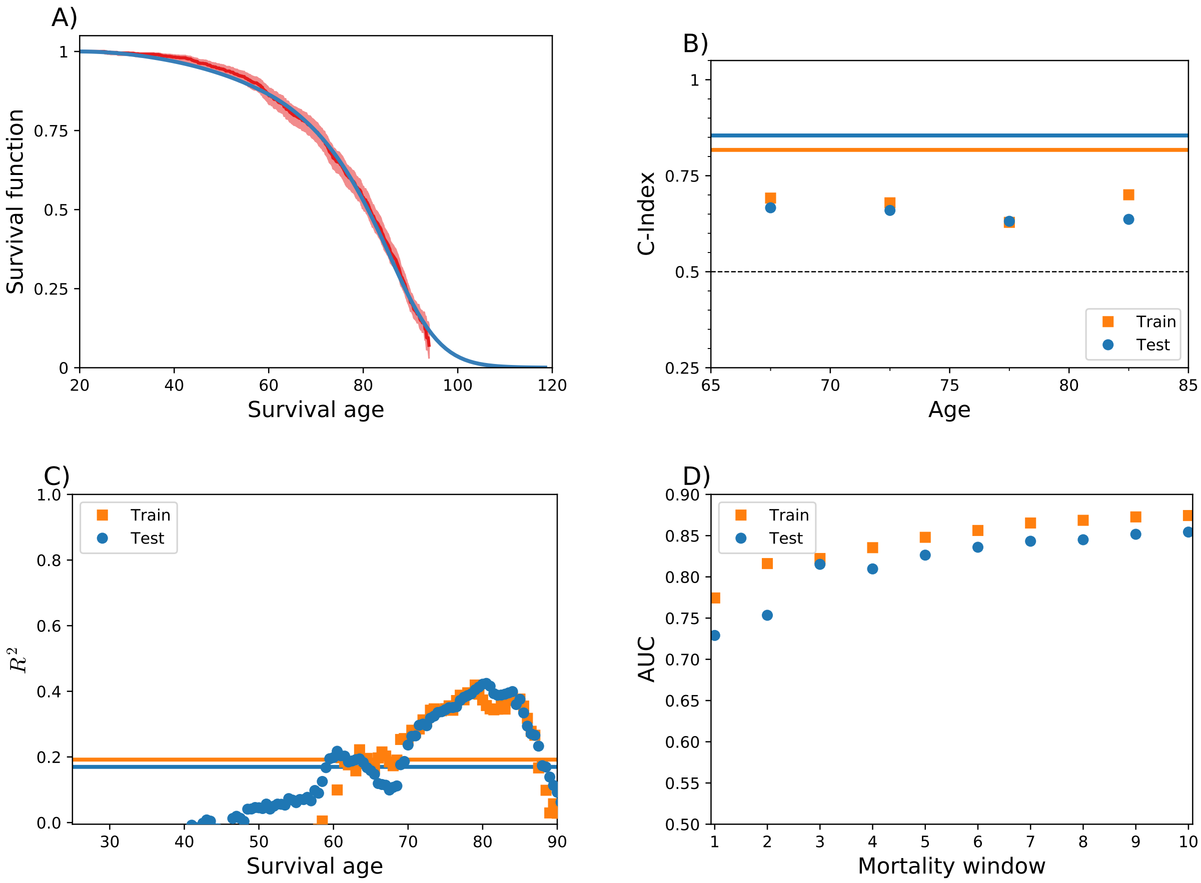


Fig S12: **NHANES survival predictions** Compare with Figs. S10 and S11. A) Model population average survival function (blue) and NHANES Kaplan-Meier population survival function (red) with a 95% confidence interval. B) Survival C-Index stratified by age (orange squares training set, blue circles test set) and unstratified (solid lines). The baseline for random predictions is $C = 0.5$ (dashed black line). C) Explained variance $R^2$ from the Brier score. Integrated $R^2$ is shown as solid lines. Higher values greater than zero mean better predicted survival curves. D) ROC AUC for the prediction of mortality within a window of time (in years) for NHANES data.

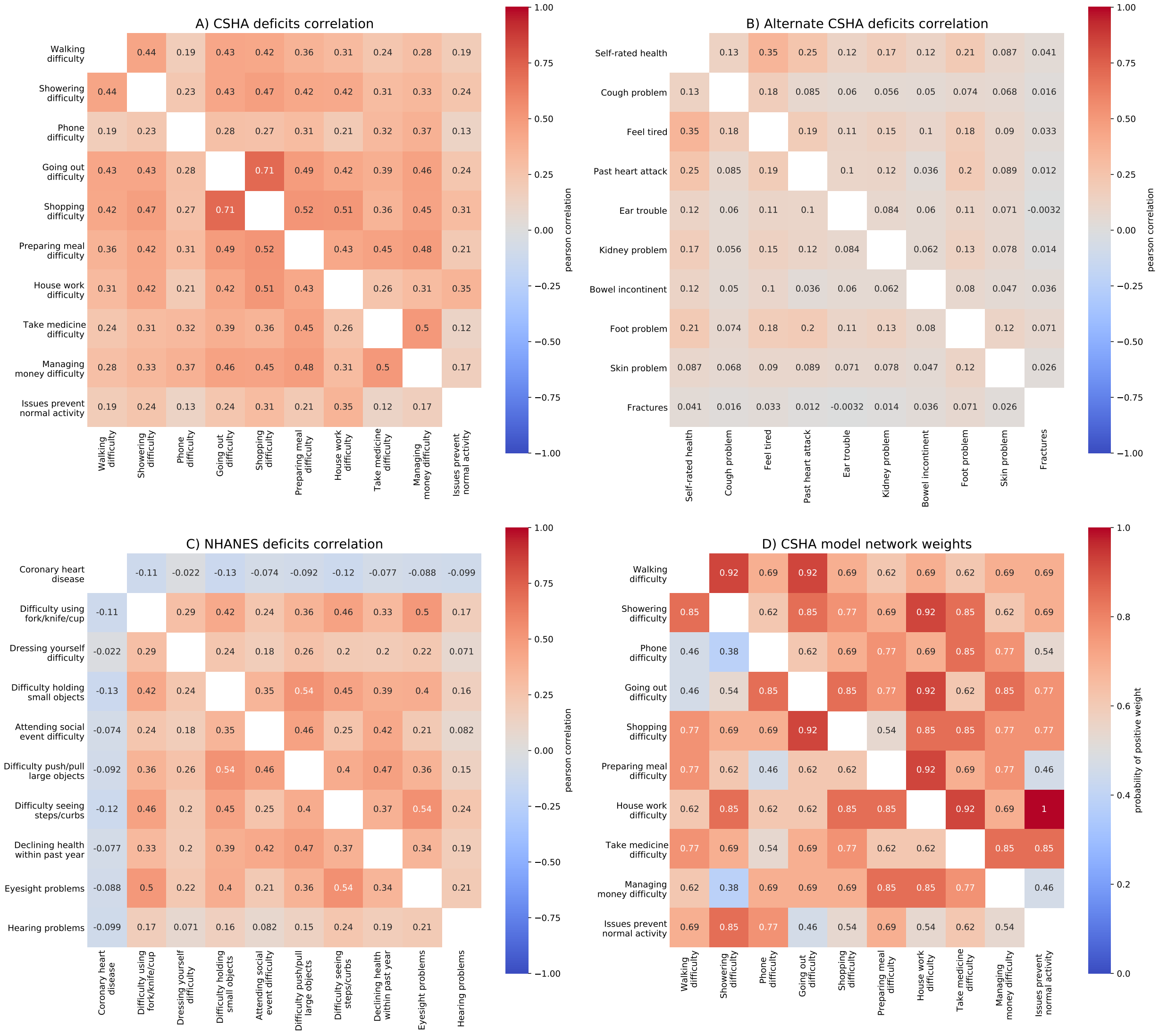


Fig S13: **Deficits correlation and network weights** A), B) and C) show pairwise correlation for the different sets of deficits used. We see that the correlation structure in all three sets of deficits is quite different. We see stronger positive correlations in the CSHA deficits used for the main plots than the alternative CSHA deficits. We generally see positive correlations for the NHANES deficits, but see a negative correlation with all deficits for the deficit "Coronary heart disease". This deficit also has slightly decreasing prevalence with age, so this is likely due to those individuals with coronary heart disease dying early. In D), we show the probability of an inferred weight in the model being positive, for 13 different model fits for the same deficits as A). The sign of the weights is quite different from the correlations. Probabilities near 0.5 mean that the inferred connection is not very robust.

### Supplemental References


[1] D. T. Gillespie. Exact stochastic simulation of coupled chemical reactions. *The Journal of Physical Chemistry*, 81:25, 1977.

[2] V. Holubec, P. Chvosta, M. Einax, and P. Maass. Attempt time monte carlo: An alternative for simulation of stochastic jump processes with time-dependent transition rates. *Europhysics Letters*, 93:40003, 2011.

[3] Michael A. Gibson and Jehoshua Bruck. Efficient exact stochastic simulation of chemical systems with many species and many channels. *Journal of Physical Chemistry A*, 104:1876 – 1889, 2000.

[4] Jeff Racine and Qi Li. Nonparametric estimation of regression functions with both categorical and continuous data. *Journal of Econometrics*, 119:99–130, 2004.

[5] Chi-Yang Chu, Daniel J. Henderson, and Christopher F. Parmeter. Plug-in bandwidth selection for kernel density estimation with discrete data. *Econometrics*, 3:199–214, 2015.

[6] James Kennedy and Russell Eberheart. Particle swarm optimization. *Proceedings of IEEE International Conference on Neural Networks*, IV(1942):1942–1948, 1995.

[7] Elre T. Oldewage, Andries P. Engelbrecht, and Christopher W. Cleghorn. Boundary constraint handling techniques for particle swarm optimization in high dimensional problem spaces. *ANTS 2018: Swarm Intelligence*, pages 333–341, 2018.

[8] Baohua Zhou, David Hofmann, Itai Pinkoviezky, Samuel J. Sober, and Ilya Nemenman. Change, long tails, and inference in a non-gaussian, bayesian theory of vocal learning in songbirds. *Proceedings of the National Academy of Sciences*, 115(38):E8358, 2018.

[9] Erika Graf, Claudia Schmoor, Will Sauerbrei, and Martin Schumacher. Assessment and comparison of prognostic classification schemes for survival data. *Statistics in Medicine*, 18:2529–2545, 1999.
